# Metabolic engineering of *Saccharomyces cerevisiae* for efficient conversions of glycerol to ethanol

**DOI:** 10.1101/2021.01.04.425180

**Authors:** Sadat M. R. Khattab, Takashi Watanabe

**Affiliations:** Research Institute for Sustainable Humanosphere RISH, Kyoto University, 611-0011, Japan; Faculty of Science, Al-Azhar University, Assiut 71524, Egypt

**Keywords:** Saccharomyces cerevisiae, bioethanol, glycerol fermentation, metabolic engineering, rewriting cofactors

## Abstract

Glycerol is an eco-friendly solvent enhancing plant-biomass decomposition through the glycell process to bio-based chemicals. Nonetheless, the lack of efficient conversion of glycerol by natural *Saccharomyces cerevisiae* restrains many biorefineries-scenarios. Here, we outline a comprehensive strategy for generating efficient glycerol fermenting *S. cerevisiae* via rewriting the oxidation of cytosolic nicotinamide adenine dinucleotide by O_2_-dependent dynamic shuttle while abolishing glycerol phosphorylation and biosynthesis pathways. By following a vigorous glycerol oxidative pathway, our engineered strain demonstrated a breakthrough in conversion efficiency (CE), reaching up to 0.49g-ethanol/g-glycerol—98% of theoretical conversion—with production rate >1 gL^−1^h^−1^ on rich-medium. Interestingly, the glycerol consumption and its fermentation unrepressed during the mixing by glucose until the strain produced >86 g/L of bioethanol with 92.8% of CE. Moreover, fine-tuning of O_2_ boosted the production rate to >2 gL^−1^h^−1^with 82% of CE. Impressively, the strategy flipped the ancestral yeast even from non-growing on glycerol, on the minimal medium, to a fermenting strain with productivities 0.25-0.5 gL^−1^h^−1^ and 84-78% of CE, respectively. Our findings promote utlising glycerol efficiently in several eco-friendly biorefinery approaches.

**Summary:** Efficient fermentation of glycerol in S. cerevisiae was established by comprehensive engineering of glycerol pathways and rewriting NADH pathway.

---

One of the challenges for sustaining the future humanosphere is the production of adequate bio-based chemicals and fuels from renewable resources with the goal of reducing greenhouse gas emissions. The paradigm of using scientific advances with metabolic engineering and biotechnology to meet the emerging needs for biofuels, as well as related materials and chemicals, has been envisioned and created on a commodity scale (Lovins *et al*, 2004; Ragauskas *et al*, 2006; Meadows *et al*, 2016). A massive requirement for ethanol recently arose for use in sanitisers because of the COVID-19 pandemic, as a concentration of 62%–71% ethanol has been demonstrated to deactivate virus particles on the skin and those that persist on inanimate surfaces such as metal, glass, and plastic (Kampf *et al*, 2020). Bakers’ yeast *(Saccharomyces cerevisiae)* has several desirable characteristics for bioethanol production, such as long and safe history of use, unicellular structure, short life cycle, remarkable fermentation abilities, robustness against inhibitors, stress tolerance during industrial levels of production, the existence of a global infrastructure for production of bioethanol from starch and molasses, and availability of tools for genetic recombination. In addition, bakers’ yeast is subject to adaptive evolution or even hybridisation and hence has been selected as a top model platform of microbial cell factories for several biotechnological applications (Peris *et al*, 2017; Khattab & Watanabe, 2019; Xiberras *et al*, 2019). The first global production of bioethanol was successfully established for blending with gasoline as a transportation biofuel. Owing to environmental, political, security and bio-economic reasons, the demand for bioethanol is increasing. However, the resources required for fermentation are limited, and challenges exist to overcome the drawbacks of second- and third-generation applications of bioethanol from lignocellulosic biomass and algae through evolving the maximum efficiencies in ethanol production from xylose and acetic acid with glucose (Luque *et al*, 2008; Deepak & Gregory, 2011; Khattab *et al*, 2013; Wei *et al*, 2013; Khattab & Kodaki, 2014).

In the past decade, glycerol-producing industries, particularly that of biodiesel, have expanded and accumulated substantial quantities of glycerol, which has led to price drops (Nomanbhay *et al*, 2018). Although the reduction degree of glycerol (C_3_H_8_O_3_) is higher than that of other fermentable sugars (Yazdani & Gonzalez, 2007), glycerol is classified as a non-fermentable carbon in native *S. cerevisiae* (Xiberras *et al*, 2019). Additionally, glycerol is a carbon source poorly utilised primarily via the glycerol-3-phosphate pathway (herein referred to as the G3P pathway), which is composed of glycerol kinase (*GUT1*) and FAD-dependent-mitochondrial glycerol-3-phosphate dehydrogenase (*GUT2*) (Sprague & Cronan, 1977). Yeast biosynthesis glycerol for mitigating osmotic stress and optimising the redox balance (Ansell *et al*, 1997). Furthermore, glycerol catabolism is subject to the repression and transcriptional regulation of glucose through respiratory factors and the *GUT1* and *GUT2* genes (Grauslund *et al*, 1999; Grauslund & Ronnow, 2000; Roberts & Hudson, 2009; Turcotte *et al*, 2010). The importance of glycerol as a carbon source that can be utilised by yeast cells has been recognised and prompted a study of the relationship between molecular inheritance and the physiology of glycerol uptake and its metabolism. The study revealed intraspecies diversity ranging from good glycerol grower to non-growers among 52 *S. cerevisiae* strains on a synthetic medium without supporting supplements and showed that the glycerol growth phenotype is a quantitative trait (Swinnen *et al*, 2013). The study confirmed that *GUT1* is one of the genetic loci that share a glycerol growth phenotype in a particular good glycerol grower, the haploid segregant CBS 6412-13A (Swinnen *et al*, 2013). Two further superior alleles of cytoplasmic ubiquitin protein ligase E3 (*UBR2*) and cytoplasmic phosphorelay intermediate osmosensor and regulator (*SSK1*) were found to link with *GUT1* for growth on synthetic medium without supporting supplements (Swinnen *et al*, 2016). These pivotal roles of *UBR2* and *GUT1* during glycerol assimilation by yeast were further confirmed by another study that resequenced the whole genomes of glycerol-evolved strains (Ochoa-Estopier *et al*, 2011; Ho *et al*, 2017). Although the G3P pathway has been evidenced as being the main catabolic pathway for glycerol catabolism in *S. cerevisiae*, its heterologous replacement with the glycerol oxidative pathway (DHA; dihydroxyacetone pathway) combined with the glycerol facilitator (*FPS1*) resulted in the restoration of growth characteristics like those of the parental strain. Furthermore, this replacement in a negative glycerol grower strain bearing the swapped UBR2_CBS6412-13A_ allele stimulated growth to the highest specific growth rate ever reported on glycerol synthetic media (Klein *et al*, 2016).

Using an approach to produce 1,2-propanediol from glycerol, a significant amount of ethanol (18 g/L) had accumulated during the first day as a byproduct, particularly on rich media (Islam *et al*, 2017). That study for production of 1,2-propanediol addressed a metabolic engineering strategy combined with heterologous replacement of the G3P route by the DHA-FPS pathway (Klein *et al*, 2016) with a module to produce 1,2-propanediol and underexpression of the triosephosphate isomerase gene (*TPI1*) (Islam *et al*, 2017). Limiting oxygen availability in shaking flask cultures resulted in increased production of ethanol from glycerol [0.165g-ethanol/g-glycerol (g^e^/g^g^) compared with 0.344g^e^/g^g^], with production rates ranged from 0.11 to 0.18 gL^−1^h^−1^on buffered synthetic medium, for facilitating the understanding the engineering of valuable products more reduced than ethanol (Aßkamp *et al*, 2019) using genetic modifications of heterologous replacement of the G3P route by the DHA-FPS pathway (Klein *et al*, 2016). It is worth emphasising that glycerol has been considered as a non-fermentable carbon source in *S. cerevisiae* (Xiberras *et al*, 2019), and attempts have been made to ferment glycerol in this yeast. These experiments were initiated by overexpressing a native DHA pathway including the glycerol dehydrogenase *ScGCY1* and dihydroxyacetone kinase (*DAK*) to produce 0.12g^e^/g^g^with a production rate of 0.025 gL^−1^h^−1^ (Yu *et al*, 2010). The methylotrophic yeast *Ogataea polymorpha* has been tested for bioethanol production from glycerol by overexpressing the genes involved in either the DHA or G3P pathways with integration of the gene for the glycerol transporter *FPS1* from *Pichia pastoris.* Furthermore, the recipient strain exhibited overexpression of the genes encoding pyruvate decarboxylase (*PDC*) 1 and alcohol dehydrogenase (*ADH*) 1. Nonetheless, overall ethanol production was relatively low (10.7 g ethanol as maximum accumulated product and 0.132g^e^/g^g^) (Semkiv *et al*, 2019). To date, there is no known native or genetically engineered yeast strain that can convert glycerol efficiently to ethanol.

Of note, we previously developed a novel biomass pretreatment method employing the glycell process with the catalysis of alum [AlK(SO4)2], with additional enhancement via microwave (Ohashi & Watanabe, 2018) There is a need for developing a *Saccharomyces cerevisiae* capable of efficiently fermenting glycerol following glycerolysis of biomass. Such glycerol-fermenting strain will co-ferment glycerol and synergist the ethanol production from hydrolysed lignocellulosic biomass. In the present study, we report the details of modelling the yeast cell to redirect glycerol traffic to bioethanol production to highest ever reported >8.6%—even in the presence of glucose— through innovation of the forthcoming systematic metabolic engineering processes outlined in Figure 1 as follows: 1) abolishment of the inherent glycerol biosynthesis pathway by knocking out NAD-dependent glycerol-3-phosphate dehydrogenase (*GPD*) 1 and retaining the second isoform of *ScGPD2* for requirements of glycerol-3-phosphate for lipid metabolism; 2) replacement of cytosolic NADH oxidation through the *ScGPD1* shuttle with a more effective O2-dependent dynamic shuttle of water forming the NADH oxidase *NoxE* to replenish NAD^+^ for the integrated gene of glycerol dehydrogenase (*OpGDH);* 3) knocking out the first gene of the G3P pathway (*GUT1*); and 4) implementing a vigorous oxidative pathway via overexpression of two copies of both the heterologous genes of glycerol dehydrogenase from *O. polymorpha (OpGDH*) and the glycerol facilitator *CuFPS1*, besides two copies of both the endogenous genes *ScTPI1* and *ScDAK1* with one copy of *ScDAK2*.

**Fig. 1.**
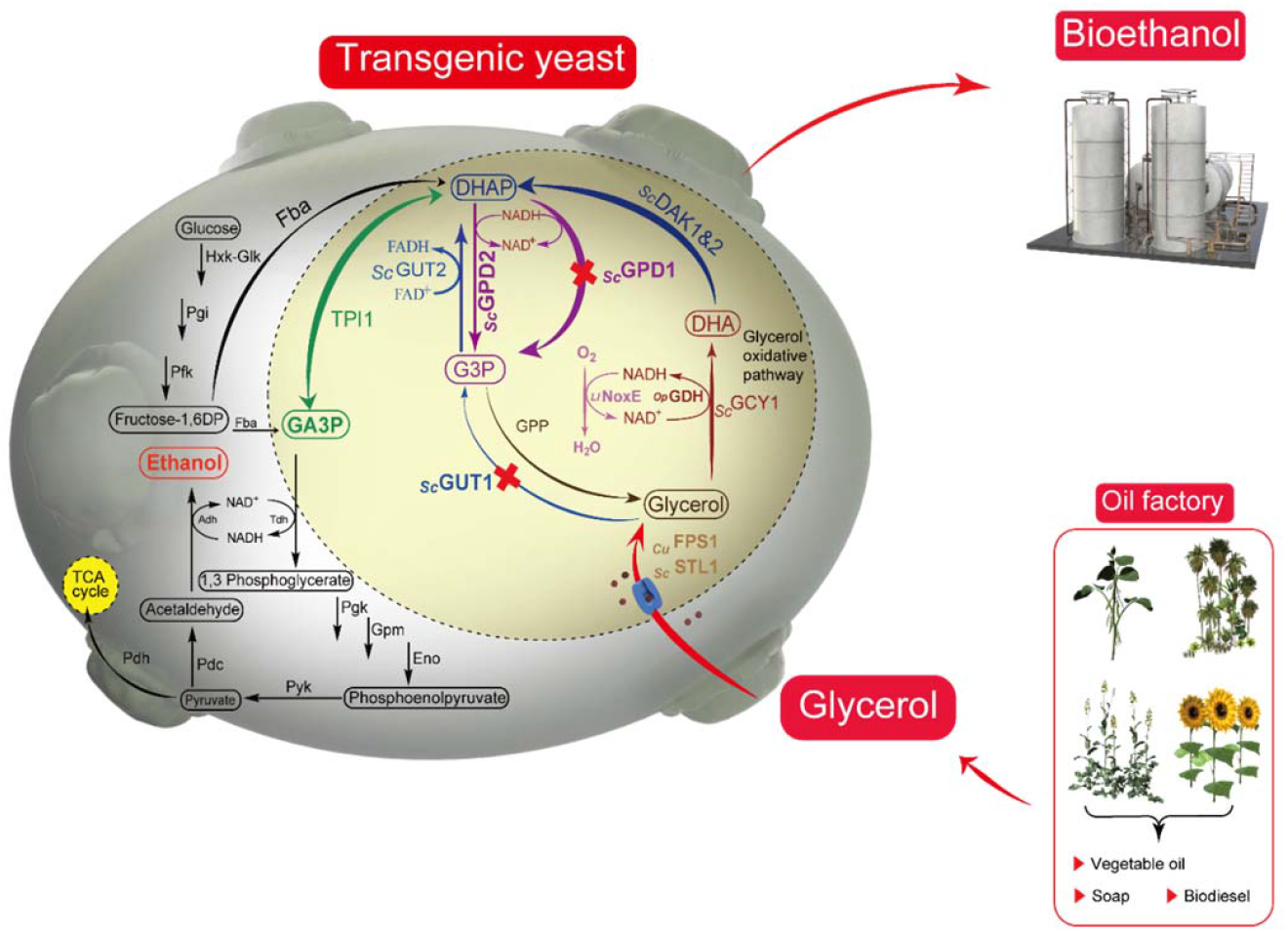
Schematic diagram showing the integrative scenario of a biorefinery with new generation of glycerol fermenting yeast and redirection of glycerol influxes to ethanol production in *Saccharomyces cerevisiae* via retrofitted native glycerol anabolic and catabolic pathways using the robust oxidative route with renovation of the NAD^+^ cofactor via O_2_-dependent dynamics of water-forming NADH oxidase. During pathway re-routing, glycerol-3-phosphate dehydrogenase 1 (*ScGPD1*) and glycerol kinase 1 (*GUT1*) were knocked-out. A highlighted circle indicates the overexpressed indigenous *S. cerevisiae* enzymes dihydroxyacetone kinase (*ScDAK*) 1 and 2 as well as triosephosphate isomerase (*ScTPI*) 1, heterologous glycerol dehydrogenase from *Ogataea polymorpha* (*OpGDH*), glycerol facilitator from *Candida utilis* (*CuFPS1*) and water-forming NADH oxidase from *Lactococcus lactis* subsp. *lactis* Il1403 (*LlNoxE*).

## Results

### Systematic metabolic engineering

#### Vigorous glycerol dehydrogenase is an essential initiator of glycerol fermentation

Initial verification of overexpression of *OpGDH* (Yamada-Onodera *et al*, 2002) in *S. cerevisiae* strain D452-2 showed strong effects compared with that of the native *ScGCY1* gene alone, or even when *ScGCY1* was integrated with other endogenous oxidative pathway genes (glycerol proton symporter of the plasma membrane *ScSTL1*, *ScDAK1*, *ScDAK2* and *ScTPI1*) in the recombinant strain GF2 (Table 1). The specific activity of *ScGCY1* in GF2 is negligible at that studied condition compared with *OpGDH* (Table 2 and the additional supplementary data). Although, the GF2 strain has a higher enzyme activity in both *ScDAK* and triosephosphate isomerase (*ScTPI1*) by 64 and 17%, respectively, compared with the strain harbouring the *GDH* gene (Table.2). As a result of the higher activity of *OpGDH*, the recombinant yeast, which was named GDH, consumed glycerol faster than GF2 in micro-aerobic condition, with a 21% increase in ethanol production, which was only 10% in GF2 as compared with that in the parental strain (Fig. 2). With semi-aerobic fermentation (10:100 medium: flask volume) using mixed glucose and glycerol for the GDH strain, improved glycerol consumption and ethanol production from 25% to 40% and 21% to 64%, respectively, before switching to the re-utilisation of ethanol when compared with previous micro-aerobic conditions (20:100 medium: flask volume) (Fig. 2 and 3). These results indicate that the first step for efficient glycerol fermentation should be through an effective *GDH* gene, initiated here with the action of *OpGDH*. Furthermore, we confirmed that glycerol consumption occurred through the constructed DHA pathway, in which its consumption was not significantly decreased after knockout of *ScGUT1*, the first gene in the G3P pathway (Fig. 2). Notably, activating the genes of the G3P pathway (i.e. *ScSTL1, ScGUT1, ScGUT2* and *ScTPI1*) in the recombinant strain GA2 (Table 1) did not result in significant improvement in the ethanol production (Fig. 2).

**Fig. 2.**
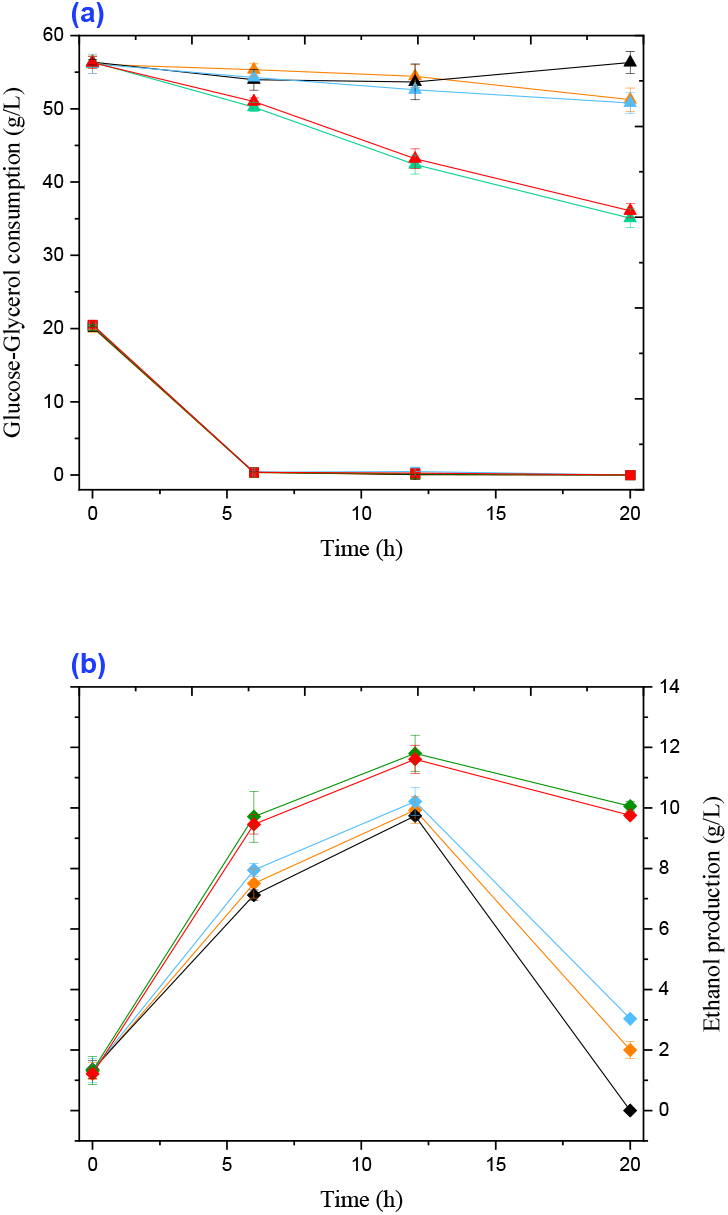
Time course for fermentation of glucose-glycerol by *S. cerevisiae* in micro-aerobic condition (20:100ml; liquid medium: flask volume) at 30°C with shaking at 180 rpm: ancestral strain (black lines); GF2 strain overexpressing endogenous oxidative (DHA) pathways *ScSTL1, ScGCY1, ScDAK1, ScDAK2* and *ScTPI1* (blue lines); GA2 strain overexpressing endogenous phosphorylation (G3P) pathways *ScSTL1, ScGUT1, ScGUT2* and *ScTPI1* (orange lines); GDH strain overexpressing *OpGDH* (green lines); ΔGUT1+ GDH (red lines); (a) glucose consumption (squares), glycerol consumption (triangles); (b) ethanol production (rhomboid symbols). Data are obtained from the mean of three independent experiments run simultaneously to decrease time differences in sampling. Error bars represent the standard deviation of the mean (n = 3). YP medium supplemented with (w/v) 2% glucose, and 5.5% glycerol was used. The only significant difference in growth rates at OD_600_ between the above strains was found on YNB medium and showed in the supplementary data (Fig. S4).

**Fig. 3.**
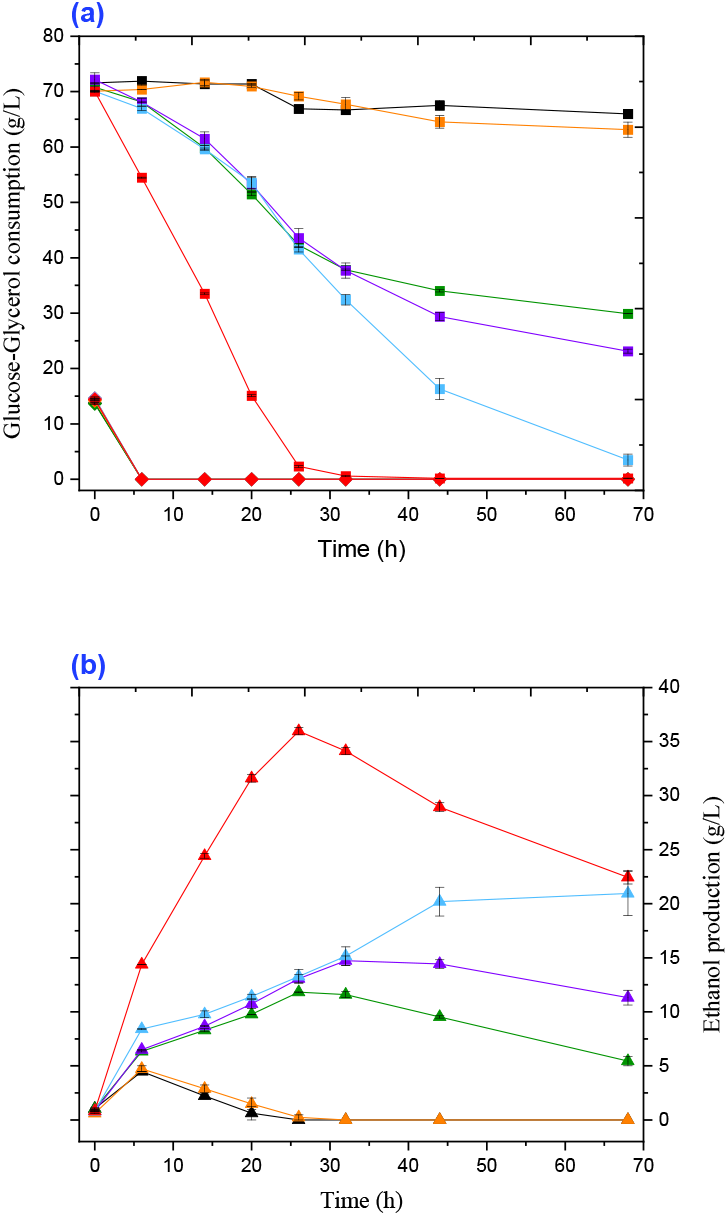
Comparison of the time course of glycerol-glucose fermentation between the *S. cerevisiae* strains within this study: ancestral strain (black lines); NoxE/*GPD1* strain (orange lines); GDH strain (green lines); GN strain (purple lines); GN-FDT strain (blue lines); GN-FDT-M1[SK-FGG] (red lines); (a) Glucose consumption (rhomboid symbols), glycerol consumption (squares); (b) ethanol production (triangles). Fermentation carried out in semi-aerobic conditions in flasks with 10:100 medium: flask volume at 30°C with shaking at 180 rpm. Data obtained from the mean of three independent experiments run simultaneously to decrease time differences of sampling. Error bars represent the standard deviation of the mean and are not visible when smaller than the symbol size. YP medium supplemented with (w/v) 1.5% glucose, and 7% glycerol was used.

**Table 1:**
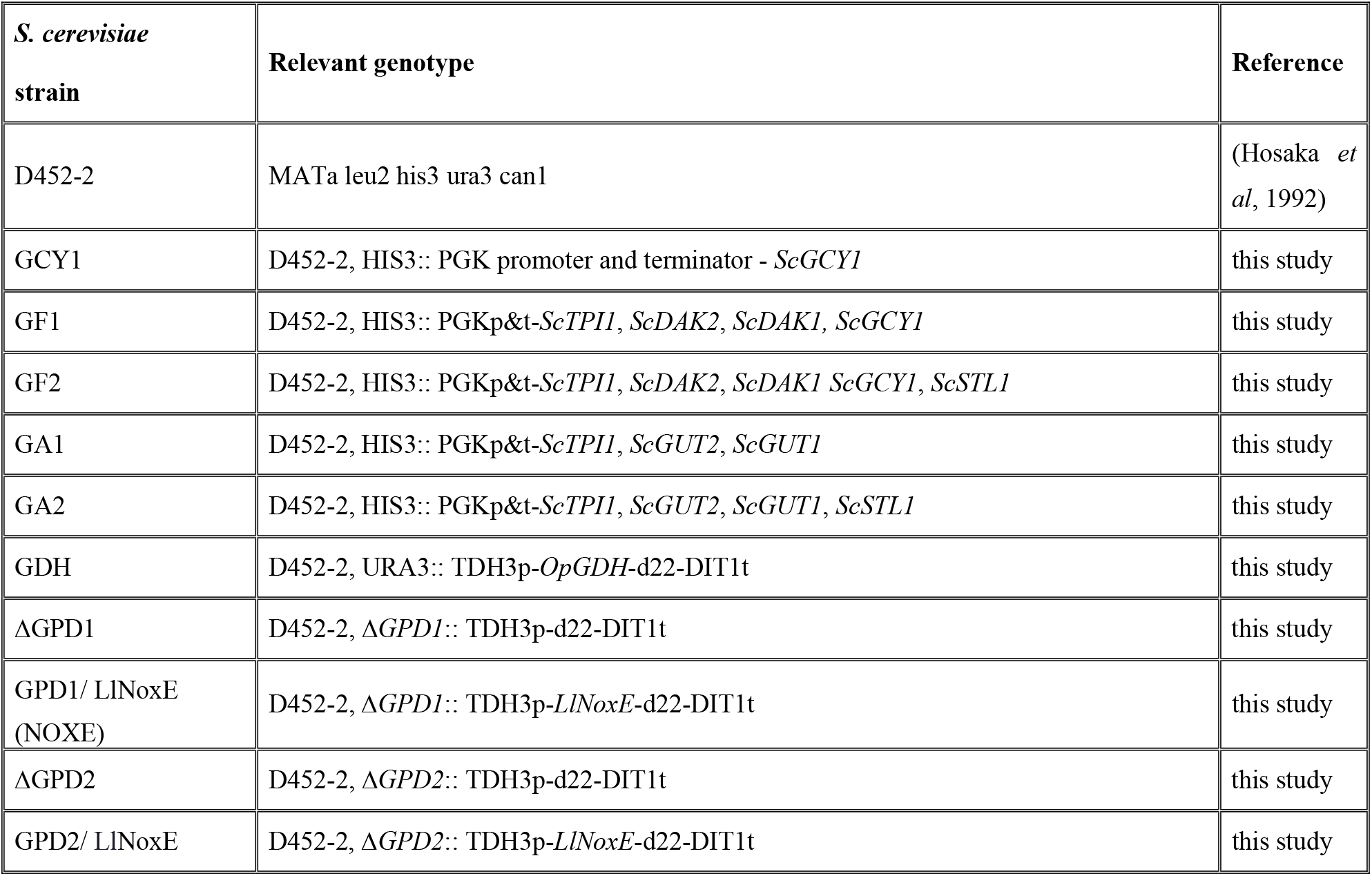
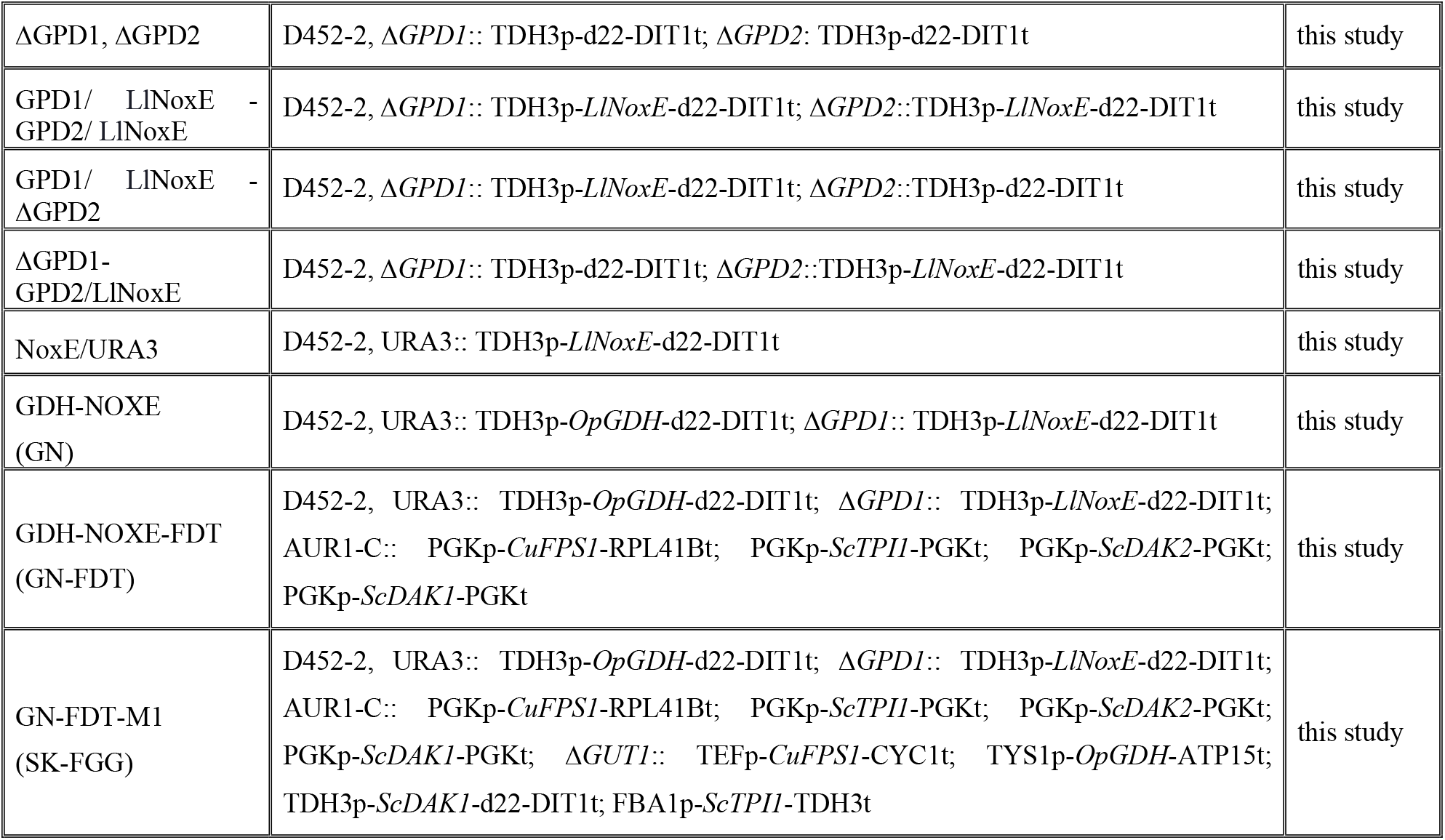
Characteristics of *S. cerevisiae* strains generated in this study:

| *S. cerevisiae*<br>strain | Relevant genotype | Reference |
| --- | --- | --- |
| D452-2 | MATa leu2 his3 ura3 can1 | (Hosaka *et al*, 1992) |
| GCY1 | D452-2, HIS3:: PGK promoter and terminator - *ScGCY1* | this study |
| GF1 | D452-2, HIS3:: PGKp&t-*ScTPI1*, *ScDAK2*, *ScDAK1*, *ScGCY1* | this study |
| GF2 | D452-2, HIS3:: PGKp&t-*ScTPI1*, *ScDAK2*, *ScDAK1* *ScGCY1*, *ScSTL1* | this study |
| GA1 | D452-2, HIS3:: PGKp&t-*ScTPI1*, *ScGUT2*, *ScGUT1* | this study |
| GA2 | D452-2, HIS3:: PGKp&t-*ScTPI1*, *ScGUT2*, *ScGUT1*, *ScSTL1* | this study |
| GDH | D452-2, URA3:: TDH3p-*OpGDH*-d22-DIT1t | this study |
| ΔGPD1 | D452-2, Δ*GPD1*:: TDH3p-d22-DIT1t | this study |
| GPD1/ LINoxE<br>(NOXE) | D452-2, Δ*GPD1*:: TDH3p-*LINoxE*-d22-DIT1t | this study |
| ΔGPD2 | D452-2, Δ*GPD2*:: TDH3p-d22-DIT1t | this study |
| GPD2/ LINoxE | D452-2, Δ*GPD2*:: TDH3p-*LINoxE*-d22-DIT1t | this study |
| ΔGPD1, ΔGPD2 | D452-2, Δ*GPD1*:: TDH3p-d22-DIT1t; Δ*GPD2*: TDH3p-d22-DIT1t | this study |
| GPD1/ LInoxE -<br>GPD2/ LInoxE | D452-2, Δ*GPD1*:: TDH3p-*LInoxE*-d22-DIT1t; Δ*GPD2*::TDH3p-*LInoxE*-d22-DIT1t | this study |
| GPD1/ LInoxE -<br>ΔGPD2 | D452-2, Δ*GPD1*:: TDH3p-*LInoxE*-d22-DIT1t; Δ*GPD2*::TDH3p-d22-DIT1t | this study |
| ΔGPD1-GPD2/LInoxE | D452-2, Δ*GPD1*:: TDH3p-d22-DIT1t; Δ*GPD2*::TDH3p-*LInoxE*-d22-DIT1t | this study |
| NoxE/URA3 | D452-2, URA3:: TDH3p-*LInoxE*-d22-DIT1t | this study |
| GDH-NOXE<br>(GN) | D452-2, URA3:: TDH3p-*OpGDH*-d22-DIT1t; Δ*GPD1*:: TDH3p-*LInoxE*-d22-DIT1t | this study |
| GDH-NOXE-FDT<br>(GN-FDT) | D452-2, URA3:: TDH3p-*OpGDH*-d22-DIT1t; Δ*GPD1*:: TDH3p-*LInoxE*-d22-DIT1t;<br>AUR1-C:: PGKp-*CuFPS1*-RPL41Bt; PGKp-*ScTPH1*-PGKt; PGKp-*ScDAK2*-PGKt;<br>PGKp-*ScDAK1*-PGKt | this study |
| GN-FDT-M1<br>(SK-FGG) | D452-2, URA3:: TDH3p-*OpGDH*-d22-DIT1t; Δ*GPD1*:: TDH3p-*LInoxE*-d22-DIT1t;<br>AUR1-C:: PGKp-*CuFPS1*-RPL41Bt; PGKp-*ScTPH1*-PGKt; PGKp-*ScDAK2*-PGKt;<br>PGKp-*ScDAK1*-PGKt; Δ*GUT1*:: TEFp-*CuFPS1*-CYC1t; TYS1p-*OpGDH*-ATP15t;<br>TDH3p-*ScDAK1*-d22-DIT1t; FBA1p-*ScTPH1*-TDH3t | this study |

**Table 2.** Specific activities of the enzymes; glycerol dehydrogenase (*GDH*), dihydroxyacetone kinase (*DAK*), triosephosphate isomerase (*TPI1*) and NADH/NAD^+^ ratio with their intracellular concentrations in the recombinant strains in this study. Error values represent standard deviation of the mean (n = 2). * indicates not detected. ¤ indicates not measured.

| Relevant strain | Enzyme activities $\mu\text{mole}/\text{min}/\text{mg}$ cell<br>extracted proteins | | | Intracellular concentrations | | NADH/ NAD<sup>+</sup><br>ratio |
| --- | --- | --- | --- | --- | --- | --- |
| | *GDH* | *DAKs* | *TPII* | NADH($\mu\text{M}$) | NAD<sup>+</sup> ($\mu\text{M}$) | |
| WT | ND* | $3.9 \pm 0.23$ | $0.058 \pm 0.003$ | $0.17 \pm 0.004$ | $0.36 \pm 0.001$ | $0.47 \pm 0.01$ |
| GF2 | ND* | $6.4 \pm 0.05$ | $0.068 \pm 0.003$ | $0.11 \pm 0.003$ | $0.32 \pm 0.002$ | $0.35 \pm 0.01$ |
| GA2 | ND* | NM□ | $0.064 \pm 0.002$ | $0.26 \pm 0.005$ | $0.67 \pm 0.008$ | $0.40 \pm 0.002$ |
| NOXE | ND* | NM□ | $0.054 \pm 0.002$ | $0.04 \pm 0.005$ | $0.36 \pm 0.004$ | $0.12 \pm 0.014$ |
| GDH | $0.063 \pm 0.001$ | NM□ | $0.055 \pm 0.003$ | $0.26 \pm 0.008$ | $0.38 \pm 0.002$ | $0.68 \pm 0.025$ |
| GN | $0.065 \pm 0.004$ | NM□ | $0.057 \pm 0.003$ | $0.34 \pm 0.018$ | $0.35 \pm 0.009$ | $0.99 \pm 0.023$ |
| GN-FDT | $0.071 \pm 0.003$ | $7.15 \pm 0.23$ | $0.067 \pm 0.004$ | $0.17 \pm 0.008$ | $0.52 \pm 0.008$ | $0.33 \pm 0.02$ |
| SK-FGG | $0.111 \pm 0.002$ | $18.3 \pm 0.28$ | $0.076 \pm 0.004$ | $0.11 \pm 0.001$ | $0.63 \pm 0.008$ | $0.18 \pm 0.005$ |

#### Efficient rewriting of the NADH oxidation pathway in *S. cerevisiae* by an O_2_-dependent dynamic water shuttle forming *LlNoxE* to replace *ScGPD1*

We comprehensively studied the replacement of *GPD* shuttles by water-forming NADH oxidase from *Lactococcus lactis* subsp. *lactis (LlNoxE).* The *GPD* shuttle is the first step in glycerol biosynthesis and represents one of the well-known systems for renovating cytosolic NAD^+^ from NADH produced during the metabolic process, such as those from the oxidation of glyceraldehyde-3-phosphate (GA3P). Consequently, glycerol is secreted upon increase of cytosolic NADH for replenishment of NAD^+^. Therefore, we first rated the participation levels of *ScGPD1* and *ScGPD2* in glycerol biosynthesis in our ancestral strain D452-2 at 10% glucose fermentation by deleting each isoform separately. In this stage, glycerol did not add to prevent interference with the biosynthesised glycerol. Besides, inducing the osmostress by the supplemented glucose at 10 % is another factor for activating *GPD* shuttles. The data revealed that the participation ratio of *ScGPD1* in glycerol biosynthesis was 82%, in which glycerol secretion from Δ*ScGPD1* was ratio compared to WT of 0.47 g/2.56 g and 23%, where 2.08 g glycerol produced from *ΔScGPD2* (Table 3; Fig. S1a). Replacing *ScGPD1* with *LlNoxE* reduced glycerol secretion by 98%, in which 0.14 g/2.56 g of WT glycerol was secreted, whereas it only reduced to 29% for the *NoxE/GPD2* strain, in which 1.82 g/2.56 g of WT was secreted (Table 3; Fig.S1a). On the other hand, replacing both *ScGPD1* and *ScGPD2* with *LlNoxE* not only prevented glycerol formation but also reduced glucose consumption significantly and obstructed cell growth and fermentation by almost the same levels (at 15%) while increasing secretion of acetic acid by 2.46-fold (Table 3; Fig.S1a-e). Similarly, replacing *ScGPD1* by *LlNoxE* while deleting *ScGPD2* had harmful effects on fermentation speed and products (Table 3; Fig. S1a-e). Comparatively, the overexpressed *LlNoxE* gene in the URA3 locus with the conserved native activity of the glycerol biosynthesis pathway exhibited a moderate reduction in glycerol production of only 41% (1.53g/2.56g) in the D452-2 strain (Table 3; Fig. S1a). Notably, replacing *GPD* with *LlNoxE* altered glycerol production to result in an increase in acetic acid production (Table 3; Fig. S1b). The efficiency of replacing *ScGPD1*, as an essential shuttle for oxidising the cytosolic NADH, by water-forming NADH oxidase was confirmed by estimating the NADH concentration, where its concentration decreased by over 4 times compared with the ancestor strain (Table 2). Exclusively replacing *ScGPD1* with *LlNoxE* is an excellent approach to eliminating glycerol formation during glucose fermentation to ethanol under the studied condition, where ethanol production increased by 9% (0.474/0.432g-ethanol /g-glucose; Table 3), therefore, consolidating this replacement with GDH strain could improve the efficiency of glycerol conversion to ethanol.

**Table 3.** Glycerol secretion, acetic acid accumulation, ethanol production, ethanol yield rate, total glucose consumed and maximum growth (measured as optical density) after 20 h of fermentation of 10%glucose as the fermentable carbon source in YP medium under Micro-aerobic conditions using recombinant strains for rewriting NADH cycles via water-forming NADH oxidase. Error values represent standard deviation of the mean (n = 3). Micro-aerobic here indicates usage of flasks with 20:100 medium: flask volume with shaking at 150 rpm. * indicates not detected.

| Relevant strain | Glycerol secreted (g) | Acetic secreted (g) | Ethanol produced (g) | Ethanol yield g/ g glucose | Consumed glucose (g) | Optical density (OD<sub>600</sub>) |
| --- | --- | --- | --- | --- | --- | --- |
| **WT** | 2.56 ± 0.06 | 1.46 ± 0.06 | 42.67 ± 0.53 | 0.431 ± 0.54 | 99.01 ± 1.660 | 13.84 ± 0.21 |
| **ΔGPD1** | 0.47 ± 0.07 | 1.66 ± 0.07 | 45.78 ± 1.09 | 0.455 ± 1.11 | 100.58 ± 2.20 | 18.10 ± 0.21 |
| **ΔGPD2** | 2.08 ± 0.05 | 1.87 ± 0.05 | 44.74 ± 0.88 | 0.446 ± 0.90 | 100.18 ± 1.10 | 17.55 ± 0.20 |
| **ΔGPD1 + ΔGPD2** | 0.10 ± 0.01 | 1.85 ± 0.03 | 43.04 ± 0.93 | 0.436 ± 0.95 | 98.81 ± 0.630 | 16.81 ± 0.17 |
| **NoxE/GPD1** | 0.14 ± 0.01 | 2.81 ± 0.08 | 47.92 ± 0.93 | 0.474 ± 0.96 | 100.97 ± 1.33 | 16.22 ± 0.19 |
| **NoxE/GPD2** | 1.82 ± 0.05 | 1.96 ± 0.06 | 44.27 ± 0.78 | 0.43 ± 0.810 | 102.34 ± 1.24 | 16.53 ± 0.27 |
| **NoxE/GPD1+ NoxE/GPD2** | ND* | 3.59 ± 0.11 | 36.24 ± 0.60 | 0.429 ± 0.62 | 84.36 ± 2.020 | 11.71 ± 0.19 |
| **NoxE/GPD1+ ΔGPD2** | 0.11 ± 0.01 | 2.60 ± 0.09 | 35.25 ± 0.61 | 0.437 ± 0.63 | 80.7 ± 2.2530 | 13.04 ± 0.18 |
| **NoxE/GPD2+ ΔGPD1** | 0.11 ± 0.02 | 2.77 ± 0.06 | 43.25 ± 0.86 | 0.435 ± 0.87 | 99.28 ± 2.110 | 13.73 ± 0.19 |
| **NoxE/URA3** | 1.53 ± 0.06 | 2.93 ± 0.06 | 43.40 ± 0.77 | 0.43 ± 0.820 | 99.7 ± 1.3230 | 18.43 ± 0.13 |

#### Integrating *GDH* and *NoxE* with *ΔScGPD1*

Based on previous results from the recombinant GDH strain and the data for replacement of oxidising shuttles of cytosolic NADH by *LlNoxE*, we studied the recycling outputs of NAD^+^/NADH between *GDH* and *LlNoxE* genes. In addition to such recycling, deleting *ScGPD1* during substitution with *LlNoxE* abolishes glycerol formation and decreases ramification of DHAP, which consolidates straightforward progression to the glycolysis route and ethanol formation. Thus, we engineered a strain that combined GDH and NOXE (Table 1). This round of recombination (GDH-NOXE or GN strain) was tested for its ability to ferment glycerol in comparison with GDH, NOXE and wild type strains. This innovative integration clearly showed improvements in both efficiency of glycerol conversion to ethanol and delay in the time of cell reprogramming to utilise the produced ethanol. After 6h both ancestral strains and the engineered NOXE strain started to re-utilise the ethanol produced from glucose without further significant consumption of glycerol, in which the maximum ethanol produced was 4.7 g/L (Fig. 3). In the GDH strain, the time for reusing the produced ethanol was delayed to 26h with increases in ethanol production to 11.82 g/L, representing 0.27g-ethanol/g-glucose-glycerol. In the case of GN strain, integration not only boosted ethanol production to 13.27 g/L (0.31g-ethanol/g-glucose-glycerol) at 26h, but also extended the fermentation time to 32h and further increased production of ethanol to 14.42 g/L before the switching to consume this ethanol (Fig. 3). In addition to the substantial activity of glycerol dehydrogenase in both strain GDH and GN, a cofactor ratio (NADH/ NAD^+^) showed a significant change from 0.68 to 0.99 between GDH and GN strains, respectively (Table. 2). This increase in the concentration of NADH is pointing to the higher efficiency of replacing *ScGPD1* by *LlNoxE* shuttle, regardless that cells have cultivated at micro-aerobic conditions (10:50 medium: falcon tubes volume).

#### Overexpressing other DHA pathway genes: *TPI1, DAK1, DAK2* and *FPS1*

Although clear impacts were observed for recycled inputs within the previous recombination, we deduced further limitations in the activity of other genes in the DHA pathway—*TPI1, DAKs* and *FPS1—* which affect the full traffic of glycerol conversion to ethanol. In this point, we must clarify that *ScDAK2* and *ScTPI1* was not considered during the previous studies of converting glycerol to ethanol. Therefore, we proceeded to overexpress the whole genes included in the DHA pathway at this stage of systematic engineering. A promoter phosphoglycerate kinase (PGK) with its terminator was used to activate the endogenous genes *ScTPI1, ScDAK1* and *ScDAK2.*The glycerol facilitator gene from *Candida utilis (CuFPS1*) (Klein *et al*, 2016) was heterologously expressed under the control of PGK promoter and the ribosomal 60S subunit protein L41B (RPL41B) terminator. The previous recombinant strain GN has been used as competent cells for receiving this one set of genes in the AUR-1C locus to generate a new strain, which was termed GN-FDT (Table 1). In the GN-FDT strain, the specific enzyme activity of *OpGDH* increased by 9% compared with the GN strain. Also*, ScDAK* and *ScTPI1* increased by 83% and 16%, respectively, compared with the WT strain. Moreover, the NADH/NAD^+^ ratio decreased to 0.33% of the GN (Table 2). This fourth recombination step (GN-FDT strain) unequivocally solved one of the main problems in this study, in which switching to ethanol utilisation was prevented before the full consumption of glycerol. The consumption rate reached 1 gL^-1^h^-1^and produced 20.95 g/L of ethanol using this recombinant strain. Nonetheless, the conversion efficiency of ethanol production appeared to be less than 48% of its theoretical value (Fig. 3).

#### Super-expressing DHA pathway using a further copy of *ScTPI1, ScDAK1, OpGDH* and *CuFPS1* genes while abolishing the native G3P pathway

The results from the fourth step of genetic engineering performed here (GN-FDT strain) highlighted the effect of the limited activities of all the genes of the DHA pathway on improving productivity with the possibility to strengthen the efficiencies by another copy of the pathway. We carefully selected and designed the strongest expression systems that were not likely to be affected by regulator repressors to constitutively express this assortment of genes (Ito *et al*, 2013; Yamanishi *et al*, 2013; Ito *et al*, 2016; Wei *et al*, 2017; Nambu-Nishida *et al*, 2018). By use of the Gibson hybrid assembly and PCR we constructed one module named M1 (Fig. S2) with the following expression systems: TEF1 promoter-CYC1 terminator, TYS1 promoter-ATP15 terminator, TDH3 promoter-mutated d22DIT1 terminator and FBA1 promoter-TDH3 terminator, for the genes *CuFPS1, OpGDH, ScDAK1* and *ScTPI1*, respectively. Therefore, we intensified an entire glycerol oxidation pathway by integrating additional copies of the genes *CuFPS1, OpGDH, ScDAK1* and *ScTPI1* during the replacement of *GUT1—* which abolished the G3P pathway—while attempting to overcome previous inadequacies during this stage of recombination. As a result, the specific enzyme activities of *OpGDH, ScDAK* and *ScTPI1* were further enhanced by 56%, 256% and 13%, respectively, compared with the GN-FDT strain. Besides, the NADH/NAD^+^ ratio decreased to 0.11 (Table 2). Interestingly, we obtained unique findings in this fourth step of recombination (SK-FGG) for glycerol consumption and ethanol production that were not previously reported in any safe organism with the developed strain GN-FDT-M1, named SK-FGG (Table 1, Fig. 3). The consumption rate of this strain reached 2.6 gL^-1^h^-1^ from glycerol at the described experimental conditions, and the productivity paced 1.38 gL^-1^h^-1^ of ethanol with a conversion efficiency of 0.44g-ethanol/g-glucose-glycerol (Fig. 3).

#### Fermentation characteristics at high initial concentrations of glycerol, with and without adding glucose, or with the higher availability of oxygen

With the current state of metabolic engineering, we examined fermentation characteristics at higher initial concentrations of glycerol (110 g/L) in the absence and presence of glucose (22.5 g/L), where glucose was reported as a suppressor for glycerol fermentation (Table 4 [A&B]). We also tested fermenting a further fed-batching of 100 g/L glycerol to the condition of [B] in (Table 4 [C]). To further confirming the engineered strain is unsubjected to repression by glucose during the glycerol fermentation, we lifted the glucose level to 45 g/L with decreased the glycerol concentration by 25% to 82 g/L before further adding 100 g/L glycerol as a fed-batch (Table 4 [D]). These outlines also testing the strain SK-FGG to produce an economically distillable ethanol titer (Table 4 [D]). The concentrations of glucose and glycerol in [D] are relatively like those obtained after the glycell process (Ohashi & Watanabe, 2018). Finally, we tested the higher oxygen availability on the production rate and efficiency at 90 g/L glycerol without glucose. The strain SK-FGG exhibited outstanding performance in micro-aerobic conditions at a higher initial concentration of glycerol–glucose mixture in YP medium, where its conversion efficiency reached 98% (0.49g^e^/g^g^) with a production rate of >1 gL^−1^h^−1^ of ethanol after consumed 82.5 g/L of glycerol from one fed-batch condition (Table 4, condition [A]). Acetic acid accumulated at 1.14 g/L at this condition (Table 4, condition [A]). Even at mixing glycerol with 22.55 g/L of glucose, its conversion efficiency was comparatively the same (Table 4, condition [B]). Interestingly, the strain engineered here is exceptional in its capacity to harmonise fermenting glycerol with glucose, along with an accumulation of >86 g/L of bioethanol with additional fed batching of glycerol (Table 4, [D]). With higher initial glucose at condition [D], cell density was promoted by 31% compared with the case [C]; besides, a minor reduction of the efficiency of ethanol conversion (Table 4, [C & D]). Notably, increasing oxygen availability by increasing flask volume by 2.5-fold while keeping broth volume constant remarkably accelerated glycerol consumption to >5 gL^−1^h^−1^ (Table 4, [E]). In addition, the rate of ethanol production increased to >2 g gL^−1^h^−1^. However, its conversion efficiency dropped to 82.8 % of the theoretical value (Table 4, [E]).

**Table 4.**
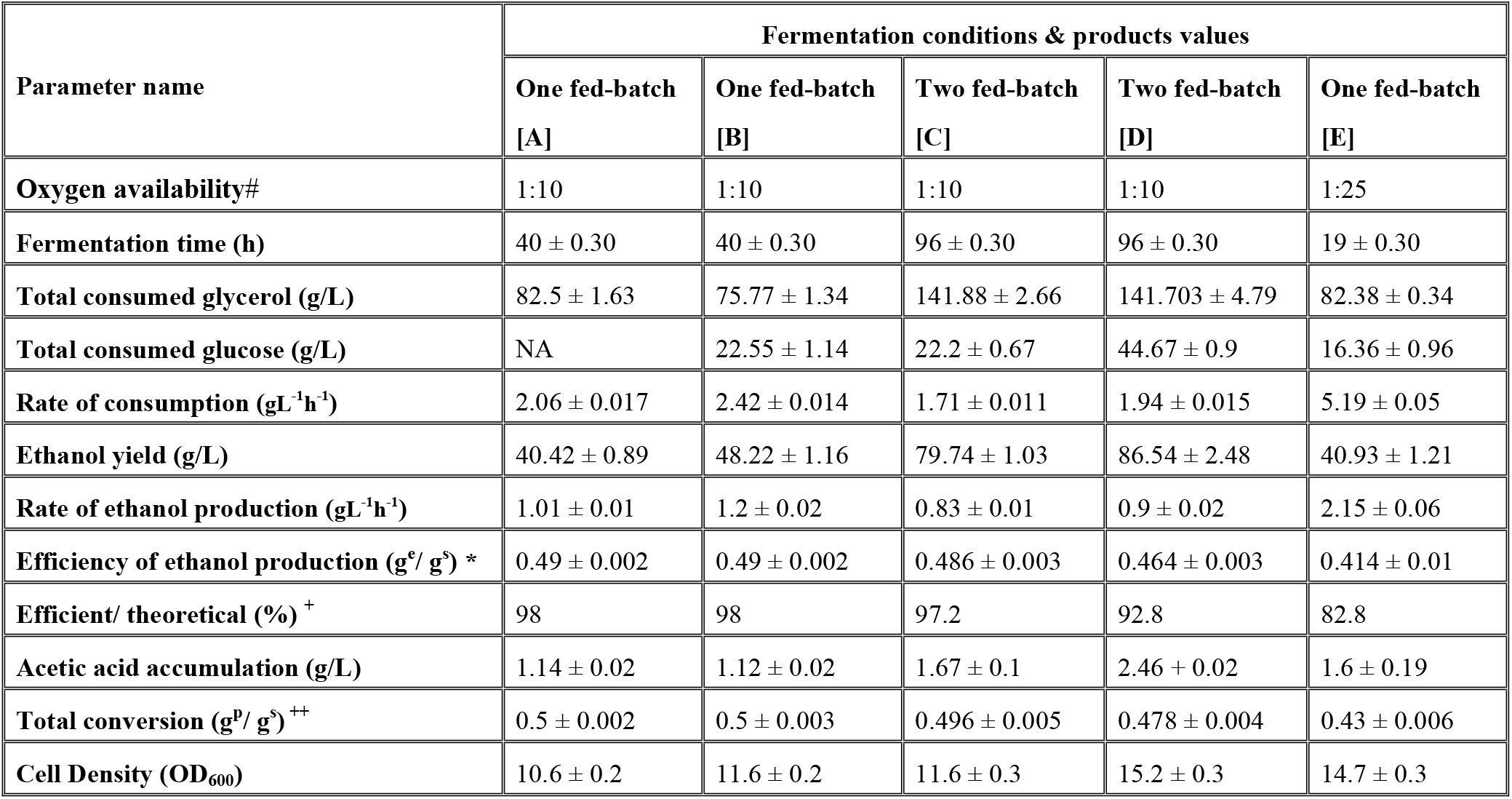
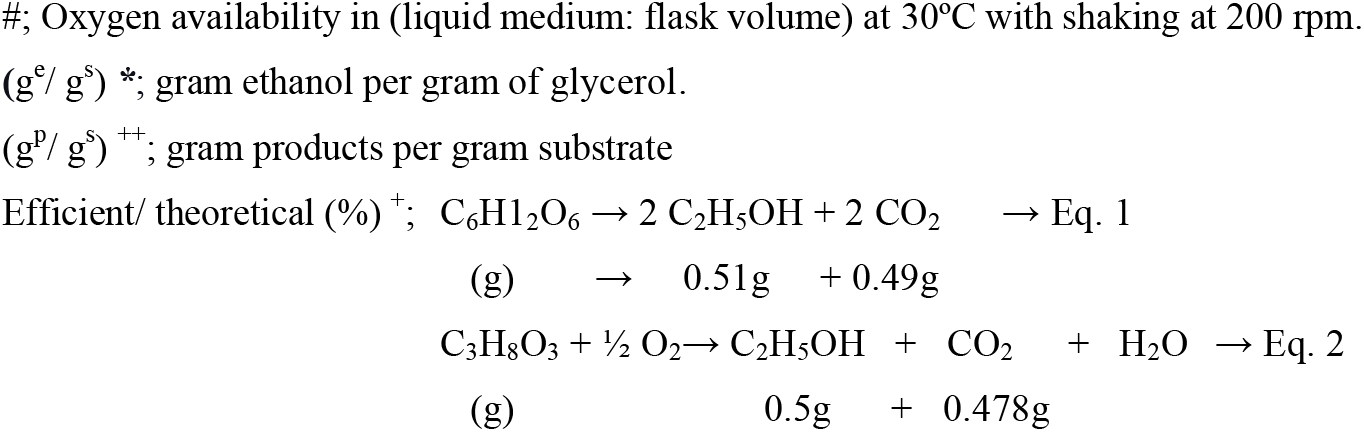
Fermentation characteristics by the generated strain SK-FGG at high concentrations of glycerol, with and without adding glucose, or with the higher availability of oxygen

#### Ferment glycerol as a sole carbon source

Our ancestor strain, D452-2 is *MATa, leu2, his3, ura3, can1* genetic base (Hosaka *et al*, 1992), and even after supplement uracil, leucine, and histidine to minimal medium, it is growthless on glycerol as a sole carbon (Fig.S4). To verify the efficiency of the best-engineered strain during this study, SK-FGG, glycerol was tested for fermentation as the sole carbon when supplemented into a yeast nitrogen base (YNB) medium with 20 mg/L of leucine and histidine. Testing was performed at four different oxygen availabilities. Under the strict anaerobic conditions, there was no further growth and no production of ethanol (Table 5). Under micro-aerobic conditions (20 ml of culture in a 100-ml Erlenmeyer flask at 30 °C with shaking at 200 rpm), 37.17 g of glycerol was consumed with a consumption rate of 0.62 gL^-1^h^-1^and a production rate of 0.25 gL^-1^h^-1^. The efficiency of ethanol conversion reached 0.42 g/g glycerol. Acetic acid accumulated at 0.78 g/L in this condition (Table 5). Glycerol was consumed more rapidly in the semi-aerobic condition (20 ml of culture in a 200-ml Erlenmeyer flask). The consumption rate was >1 gL^-1^h^-1^, which raised the ethanol production rate to 0.44 gL^-1^h^-1^. There was 2.88 g of acetic acid cumulation in this condition. As a result, the total conversion reached 0.44 g/g glycerol (Table 5). With further elevation of the oxygen availability in 20:300-ml liquid medium: flask volume ratio, the rate of glycerol consumption and ethanol production rate was boosted to 1.29 and 0.5 gL^-1^h^-1^, respectively. The efficiency of ethanol conversion was 0.39g^e^/g^g^, and the total convertibility was 0.45g^e^/g^g^, which represents 90% of the theoretical conversion regardless of the utilised glycerol in cell formation (Table 5).

**Table 5.**
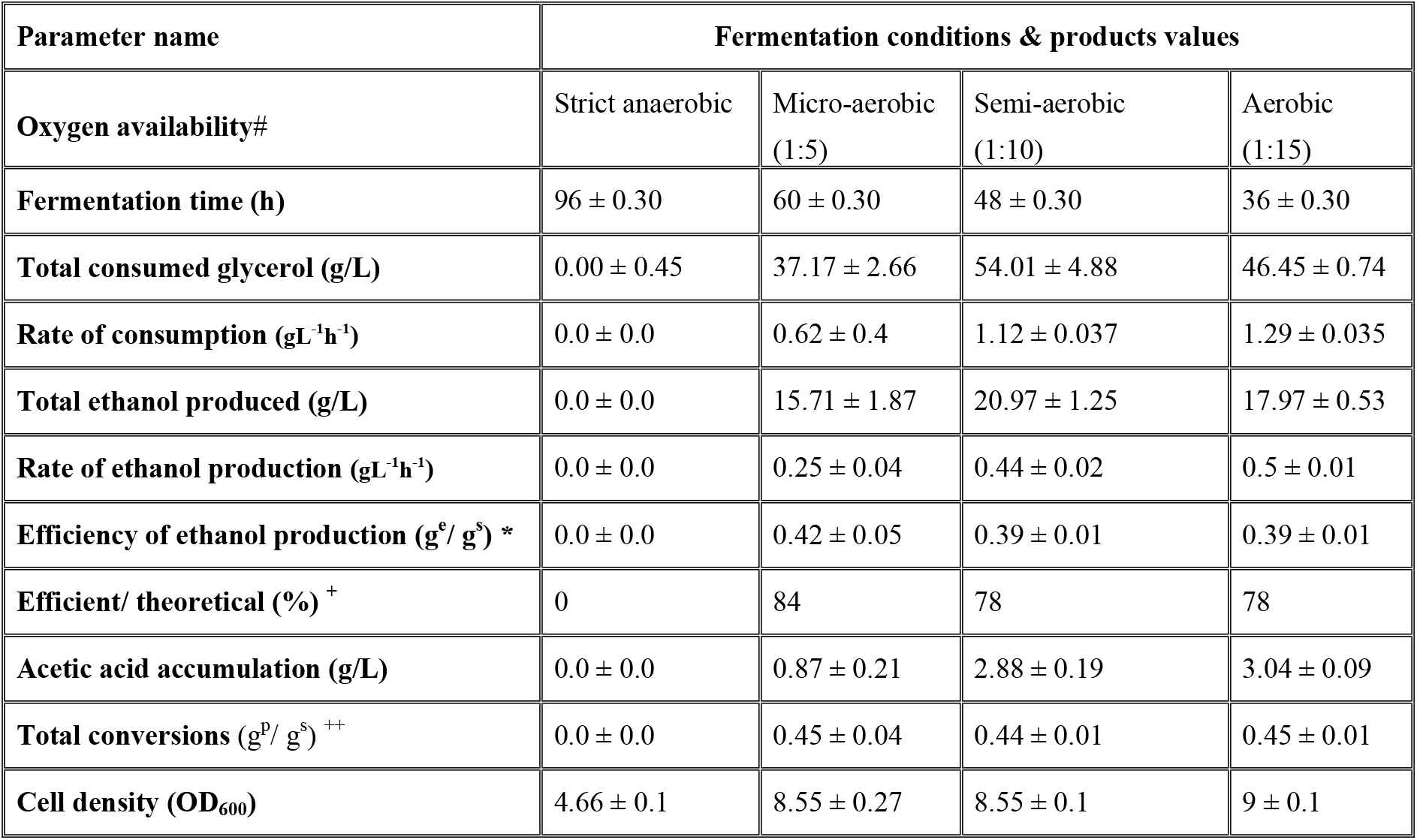
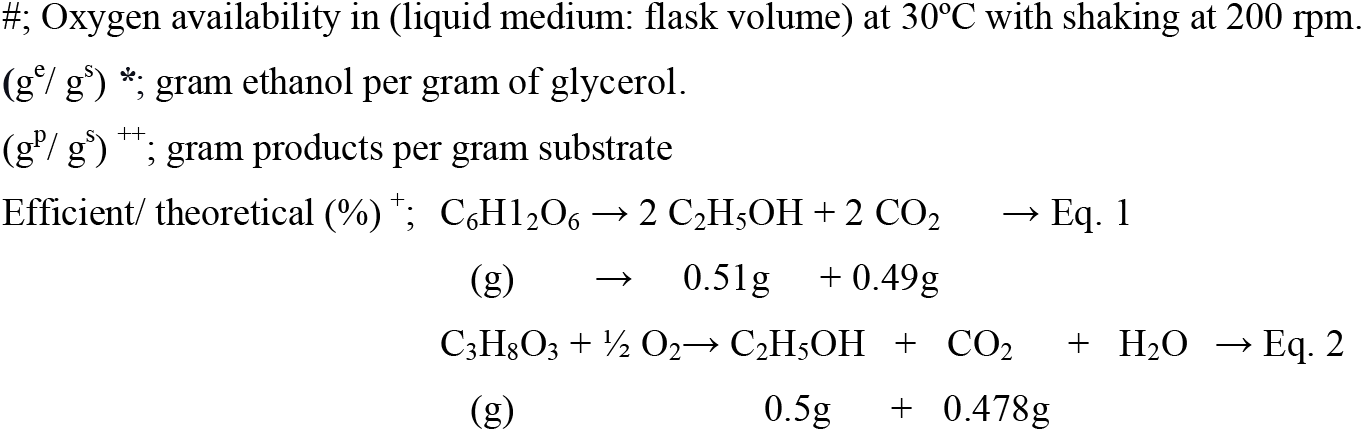
Fermentation characteristics of glycerol as a sole carbon source by the generated strain SK-FGG at different oxygen availabilities.

## Discussion

Recently, microbial technologies for exploiting glycerol as a carbon source for producing valuable products have gained increasing attention, where a considerable amount of glycerol accumulates as an unavoidable byproduct from the expansion of biodiesel industries. Moreover, we have developed a novel promising pretreatment method using glycerol for support the solvolesis of lignocellulosic biomass with alum and microwaving (Ohashi & Watanabe, 2018). Therefore, generating a yeast strain capable of converting such a mixture of glucose and xylose with glycerol following glycerolysis was inevitable for developing our methodology. Here we describe the initial study to construct a novel strain that can efficiently convert pure glycerol with glucose to ethanol.

It is well known that native *S. cerevisiae* contains full genes for two metabolic pathways (DHA and G3P) for glycerol catabolism (Fig. 1); nonetheless, glycerol was considered as non-fermentable and an unfavourable carbon source as a feedstock (Sprague & Cronan 1977; Xiberras *et al*, 2019). Furthermore, one of the ancestral of our laboratory strain belongs to S288C (Kodaki & Yamashita 1989; Hosaka *et al*, 1990; Hosaka *et al, 1992*) (Table 1 and Fig. S4) that has been classified as a negative glycerol grower on synthetic medium (Swinnen *et al*, 2013). Therefore, we tested our strain D452-2 for growth on glycerol as a sole carbon source in YNB medium supplemented by 20 mg/L of leucine, histidine and 5 mg/L of uracil. The maximum optical density obtained after the growing for 180h was (0.28). That data evidences the classification of the ancestral D452-2 strain in a negative glycerol grower class (Fig. S4). Hence, herein we initially focused on using a conventional YP medium supplemented glycerol or/and glucose to overcome the limiting nutritional factors that arise from using a synthetic or minimum medium, as well as the necessity for providing amino acids and nucleic bases. Moreover, this YP medium showed enhancement in bioethanol accumulation from glycerol during the onset of 1, 2-propanediol production (Islam *et al*, 2017).

Although abolishing *ScTPI1* has been a pivotal hub to produce glycerol from glucose (Overkamp et al, 2002), which was reversed here, this gene has not been integrated with a previous study examining overexpressing the native DHA-pathway (Yu *et al*, 2010). Therefore, we initiated our study by updating that work (Yu *et al*, 2010) via combination of overexpression of *ScTPI1* and *ScDAK2* with the DHA pathway and replaced a previously reported *GUP* gene with a currently unique and generally accepted glycerol transporter *ScSTL1* to track restrictions in the native oxidative pathway during glycerol fermentation. Hence, we constructed the strain GF2, which overexpressed the genes *ScSTL1*, *ScGCY1*, *ScDAK1*, *ScDAK2* and *ScTPI1*. Additionally, we constructed the GCY strain, which overexpressed only the *Sc*GCY1 gene and GF1, which activated the genes *ScGCY1, ScDAK1, ScDAK2.* Similarly, we generated GA1, which overexpressed *ScGUT1*, *ScGUT2* and *ScTPI1*, as well as GA2, which activated *ScSTL1*, *ScGUT1*, *ScGUT2* and *ScTPI1*.

Albeit we detected a significantly faster growth rate, particularly for GF2 (Fig. S4), we did not observe a significant improvement in ethanol production with that strain overexpressed its *ScGCY1* only at the conditions tested in Fig. 2 (data omitted to avoid overlap with the ancestral strain). However, we detected significant ethanol production by GA2 and GF2 that was rapidly re-consumed in the presence of higher concentrations of glycerol (Fig. 2). Concurrently, we recognised the limited activity of the endogenously overexpressed *ScGCY1* gene compared with glycerol dehydrogenases from *O. polymorpha* (Nguyen & Nevoigt, 2009). Therefore, we constructed a yeast harbouring *OpGDH* named GDH to be tested with GF2 during glycerol fermentation. Furthermore, we confirmed the limited activity of the overexpressed *ScGCY1* gene compared with glycerol dehydrogenases from *O. polymorpha* (Table 2). As a result of these comparative studies, such an active *OpGDH* gene is the first key for deciphering glycerol fermentation, although the sole integration of *OpGDH* was not enough to induce an efficient conversion (Fig. 2). On the other hand, overexpressing the native glycerol catabolic pathway G3P in the GA2 strain did not demonstrate promising results compared with this oxidative pathway. We hypothesise that this is a limit of the respiratory chain during glycerol consumption and thus, restricted the renovation of FAD^+^ for converting glycerol-3-phosphate to DHAP through *GUT2*, considering that this phosphorylated G3P pathway is subject to repression and transcriptional regulation with the presence of glucose (Grauslund *et al*, 1999; Grauslund & Ronnow, 2000; Roberts & Hudson, 2009; Turcotte *et al*, 2010; Saito & Posas, 2012; Babazadeh et al, 2017).

Besides the induction for expressing target genes, one of the other main obstacles affecting the efficiency of microbial production is to meet the stoichiometries of the engineered metabolic pathways, cofactors and ATP/oxygen ratios, especially for those pathways which require cofactors for their activation (Vemuri *et al*, 2007; Deepak & Gregory, 2011; Meadows *et al*, 2016). As an integrated potent *OpGDH* in *S. cerevisiae*,we visualised that *GPDs* shuttles may promote activation of oxidation of the plethora from cytosolic NADH (Table 2) and a reduction of DHAP into glycerol biosynthesis pathway. Additionally, a ramification of glycerol-3-phosphate into glycerolipid pathway may take place (Zheng & Zou, 2001). Likewise, DHAP may be distributed into phospholipid and methylglyoxal biosynthesis (Murata *et al*, 1985; Zheng & Zou, 2001). On the other hand, external mitochondrial NADH dehydrogenase (*NDE*) 1 showed involvement in recycling the cofactor during glycerol metabolism (Aßkamp *et al*, 2019). Furthermore, it is well known that the alcohol dehydrogenase *ADH2* primarily catalyses the reaction of oxidising ethanol to acetaldehyde and the synthesis of *ADH2* is repressing by glucose and vice versa in the presence of glycerol (Lutstorf *et al*, 1968). The activity of *ADH* within cells grown in glycerol yeast extract medium has been shown to be ten times higher as compared to those cultured in glucose (Lutstorf *et al*, 1968). Further investigations are needed to elucidate and quantify the participation of each of the five isoforms of *ADH* during switching and utilise an ethanol produced concomitant the glycerol.

In native yeast with access to glucose, especially in the presence of oxygen, there is a plethora of cytosolic NADH. As a result, there is a need for shuttles to re-oxidise this surplus. The *GPD* shuttle plays an essential role in this regard with reduction of DHAP to glycerol-3-phosphate to maintain this homeostasis (Larsson *et al*, 1998). Intracellular redox homeostasis in *S. cerevisiae* comprises > 200 reactions; thus, shuttles oxidising NADH have been well studied (Ansell *et al*, 1997; Vemuri *et al*, 2007). One successful strategy involves catalysing the oxidation of cytosolic NADH by heterologous expression of a water-forming oxidase gene in *S. cerevisiae* for reducing the cytosolic NADH and revert the glycerol biosynthesis from glucose to 2, 3-butanediol, and acetoin (Kim *et al*, 2015; Kim & Hahn 2015; Bae et al, 2016; Kim *et al*, 2019). Also, the combination uses *NoxE* and *GPDs* during the conversion of glucose to ethanol has been reported (Vemuri *et al*, 2007). Nonetheless, there are no studies regarding the effects of replacing the native shuttles of *GPD* with these alternative shuttles (*NoxE*) for oxidising cytosolic NADH during ethanol production from glucose. Also, there are no reports about using *NoxE* during the glycerol conversion to bioethanol. Therefore, in the second round of recombination described here, we comprehensively focused on preventing that overflow to glycerol biosynthesis with the conservation of intracellular redox homeostasis during glucose fermentation under micro-aerobic condition. Investigation of the nine constructed strains of either deleted or replaced *ScGPD1* and/or *ScGPD2* by the *LlNoxE* gene indicated that replacing *ScGPD1* by *LlNoxE* is the best approach, in which glycerol biosynthesis was effectively abolished by 98%, and improvement in fermentation efficiency by 9% was observed (Table 3). This strategy remarkably decreased cellular NADH concentration by 76% under limited the oxygen availability (Table 2).

Expectedly, this single gene replacement will not exhibit significant progress toward glycerol fermentation (Fig. 3). The low activity of native glycerol dehydrogenase *ScGCY1* is known (Nguyen & Nevoigt, 2009). We confirmed there is no detectable activity in our strain under that investigated conditions (Table 2). As detailed above, ramification of DHAP represents another hindrance for straightforward progress into the glycolysis route from glycerol. At this juncture, reduced circulation of DHAP into the glycerol biosynthesis and G3P pathways was confirmed to be efficient for glycerol fermentation by integrating this replacement of *ScGPD1* by *LlNoxE* within the GDH strain. As expected, the strain harboured this unique point of integration (GDH-NOXE) showed substantial improvement in ethanol production from glycerol, which reached 28% compared with the GDH strain at the studied conditions, and was not considered for other parameters, such as oxygen levels (Fig. 3). The role of abolishing *ScGPD1* was explicitly calculated from the data (Fig. 3), which represented 43% of that improved ratio. The balance of cofactor was confirmed in this combination by measuring the cellular content, as shown in Table 2. Utilising the recycled cofactors of NADH/NAD^+^for production of 1, 2-propanediol has been well studied during glycerol fermentation (Islam *et al*, 2017).

The importance of activation of the other genes in the DHA pathway has been confirmed through continued bioethanol production until full glycerol consumption (Fig. 3). Although we did not evaluate the effect of overexpressing each gene individually, we estimated the specific activities of *OpGDH, ScDAKs, ScTPI1* in the recombinant strain and recognised the cooperative effects for overcoming that traditional and ambiguous phenomenon of re-consuming the produced ethanol earlier than the occurrence of the full consumption of glycerol. As projected, an increase in the activity of *ScDAKs* plays another essential role with both *OpGDH* and *ScTPI1* for glycerol conversion, where GN-FDT showed higher specific enzyme activities of *OpGDH, ScDAKs*, and *ScTPI1* compared with GN and the ancestor strain (Table 2). Moreover, conservation of the NADH/NAD^+^ at the lower ratios is substantial for the straightforward reaction toward the ethanol formation. In this regard, it was reported that the permeability of the three-carbon compounds, including glycerol in *Candida utilis*, is much more rapid than that in bakers’ yeast, which supports the efficient utilisation of glycerol even at low concentrations (Gancedo *et al*, 1968). Therefore, we heterologously expressed *CuFPS1* in *S. cerevisiae* to support the influxes of glycerol in our strain as also reported previously (Klein *et al*, 2016; Islam *et al*, 2017). Besides, *ScDAK1* and *ScDAK2* were characterised for detoxifying DHA, with *K*m(⊓HA) of 22 and 5 μM and *K*m(_ATp_) of 0.5 and 0.1 mM, respectively (Molin *et al*, 2003), thus overexpressing *ScDAK2* (which has a much lower *K*m_(DHA-ATP)_) with *ScDAK1* definitely detoxified the DHA that may accumulate by the action of the introduced *OpGDH* and *CuFPS1* in this study. Deletion of *DAK*s resulted in the accumulation of significant amounts of DHA during glucose fermentation (Nguyen & Nevoigt, 2009). In addition to detoxification, overexpressing *DAKs* was thought to speed up the transfer of DHA to DHAP, where overexpressed *DAK*1 reduces DHA accumulation by nearly 16% compared with that of the overexpressed strain *OpGDH* (Klein *et al*, 2016). Although we tested the extracellular accumulation of both DHA and DHAP in our recombinant strains, we found that their concentrations were below detectable levels. This may be because our ancestor strains have *DAKs* activity at 3.9 μmole/min/mg of cell extract protein (Table 2). Nevertheless, with the genetic modifications presented during the introduction of *CuFPS1* and *OpGDH*,while overexpressing *ScDAK1* and *ScDAK2*, DHAP may have intracellularly concentrations of influxes into the glycerol biosynthesis and G3P-pathways through *ScGPD2* or to saturate the native activity of *ScTPI1* to convert into the pentose phosphate pathway, especially with the presence of glucose (Grüning *et al*, 2014). Through scrutiny of the previous studies in which the activity of *ScTPI1* was abolished, we recognised the pivotal role of overexpressing *ScTPI1* in the current study, where the intracellular concentration of DHAP accumulated to 30-fold concentrations (Shi *et al*, 2005) and when this deactivation was further coupled with additional deletions of *ScNDE1*, *ScNDE2* and *ScGUT2*, the fermentation product shifted from ethanol to glycerol (Overkamp *et al*, 2002).

However, integrating one copy of the whole DHA pathway with *LlNoxE* generated the ability in yeast to convert all supplemented glucose and glycerol to ethanol. Nonetheless, we recognised that the conversion efficiency may still be affected by the robustness of native programmed glycolysis. Besides, the availability of ATP for *DAKs* was proposed as another limiting factor because DHA accumulation was decreased by nearly 58% with replacing a phosphorylation pathway through *GUT1* by a synthetic DHA pathway (Klein *et al*, 2016). Hence, this obstacle could be overcome by further strengthening of the entirety of the genes in the DHA pathway via additional copy under different expression systems with the replacement of *GUT1*. Potentially employing the strategy of multi-copy gene integrations while optimising the stoichiometries of the metabolic pathway considerably boosted production, e.g., six copies of the farnesene synthase gene, which was integrated into yeast to improve the synthesis of farnesene (Meadows *et al*, 2016). Here, with integration of a second copy of genes *CuFBS1*, *OpGDH*,*ScDAK1* and *ScTPI1*, we further selected highly constitutive expressing systems in yeast (Ito *et al*, 2013; Yamanishi *et al*, 2013; Ito *et al*, 2016; Wei *et al*, 2017; Nambu-Nishida *et al*, 2018) to extend production levels and efficiencies, using the TEF1 promoter-CYC1 terminator, TYS1 promoter-ATP15 terminator, TDH3 promoter-mutated d22DIT1 terminator, and FBA1 promoter-TDH3 terminator, respectively. The other copies of *CuFPS1, OpGDH, ScDAK1* and *ScTPI1*, which replaced *ScGUT1*, strengthened the specific activities of enzymes *OpGDH, ScDAKs*,and *ScTPI1* by further 56%, 256% and 13%, respectively, compared with the GN-FDT strain. Also, the NADH/NAD^+^ ratio decreased to 0.11 (Table 2). Interestingly, these reinforced activities with the possibility of providing ATP from the deletion of the *GUT1*to the *DAKs* boosted production rates from 0.85~1.28 gL^−1^h^−1^ and the conversion efficiency reached 98% of the theoretical ratio in the rich medium (Table 4 [A]). Furthermore, the strain overrode the glucose repression during glycerol fermentation with glucose and produced up to 8.6% of ethanol (Table 4 [D]). For the efficient SK-FGG strain generated and its introduced pathway, oxygen availability became the limiting factor. Surprisingly, fermentation rates doubled with the fine-tuning to >2 gL^−1^h^−1^ (Table 4 [E]) although the CE decreased to 82.8% of the theoretical conversion.

Interestingly, our strain SK-FGG showed abilities to convert glycerol, as a sole carbon source supplemented in YNB medium (without supplementary amino acids), to ethanol with production rates of 0.25 and 0.44 gL^−1^h^−1^ with efficiencies 0.42 and 0.39g^e^/g^g^, respectively with micro and semi-aerobic conditions (Table 5). Also, fine-tuning the oxygen availability shows notable effects during our study by decreasing the flasks sizes where it boosted the conversion efficiency from 78% to 84%. This result matches with the previous finding, which revealed an improvement in ethanol production from 0.165g^e^/g^g^ to 0.344g^e^/g^g^ by reducing the oxygen availability during fermentation of glycerol in a buffered synthetic medium (Aßkamp *et al*, 2019). Later, *NDE* genes have reported as the main shuttles for re-oxidising cytosolic NADH during the growth and conversion of glycerol after replacing *GUT1* with a DHA-FPS pathway in best natural positive-growing strain on synthetic medium supplemented by glycerol (Swinnen *et al*, 2013; Aßkamp *et al*, 2019). As a result, the conversion rates and maximum titer productivities are bounding by the oxidising shuttles within the respiratory chain and the availability of the final electron acceptor (Aßkamp *et al*, 2019). In addition to the competition with cell biomass formation (Aßkamp *et al*, 2019). As a result, the ethanol productivity rate was limited between 0.11~0.18g gL^−1^h^−1^, although higher cell biomass levels (Aßkamp *et al*, 2019). Moreover, the highest ethanol titer was 15.7 g/L.

On the other hand, renovate cytosolic NADH by replaces *ScGPD1* with *LlNoxE* showed higher potentials in productivity rates with higher efficiencies. Even when using glycerol as a sole carbon source in an unbuffered minimal medium, our engineered strain SK-FGG showed higher production rates ranged from 0.25 to 0.5 gL^−1^h^−1^ with higher conversion efficiencies set between 0.42 to 0.39g^e^/g^g^, respectively. It is evidenced that SK-FGG fermented glycerol without linking with the respiratory routes and its limitation or the competition with cells formation, where SK-FGG reached the maximum growth of 8.55~9 OD at 12h and continued glycerol consumption and ethanol production until reaching >22 g/L without further cell growth (supplementary data for Table 5). By considering the start point of the ancestor strain, which showed growthless on glycerol as a sole carbon source (Fig.S4) until reached by growth levels to 8.55 or 9 OD in SK-FGG (Table 5). With the productivity rates in this study, re-cycling NADH by *LlNoxE* during replacing *ScGPD1*, which concomitant abolishing glycerol biosynthesis, is an innovative point for fermenting glycerol and supported the conversion to unprecedented production rates and efficiencies. Besides, the other novelty of overexpressing *ScTPI1* and *ScDAK2*during glycerol or/and glucose fermentation.

The remarkable difference between YP and YNB media is the increased shift from ethanol production to acetic acid accumulation of 2.88g from 54.01g of consumed glycerol in semi-aerobic conditions. As a result, the total conversion of acetic acid and ethanol reached 90% of the theoretical value (Table 5). Like YP, a fully aerobic condition promoted the acceleration of fermentation with the same efficiency of ethanol formation. The content of amino acids in the medium has a crucial role in cell growth (Roberts et al, 2020); furthermore, we observed *de novo* biosynthesis of NAD^+^ from tryptophan through the kynurenine pathway (Panozzo *et al*, 2002; Kato & Lin, 2014). Nicotinic acid, nicotinamide, quinolinic acid and nicotinamide riboside can salvage the NAD^+^ biosynthesis (Kato & Lin, 2014). In this regard, nicotinic acid is auxotrophic under anaerobic conditions in *S. cerevisiae* (Panozzo *et al*, 2002). These limitations may explain the slower conversions and lower efficiency of ethanol production from glycerol when using YNB (without supplemented amino acids) compared when using YP medium. We must emphasise that strain SK-FGG was unable to ferment glycerol under the strictly anaerobic condition in our experiments, which used YNB medium (Table 5). Undoubtedly, there are no shuttles for the renovation of NADH under anaerobic conditions. This finding agrees with the previously published data on the effect of oxygen limitation on preventing the production of DHA during the fermentation of glucose (Nguyen & Nevoigt, 2009). Therefore, this study shows that the recycling of cofactors is sufficient for complementing the robust oxidation of the DHA pathway for efficient utilisation of glycerol to produce bioethanol or other bio-based chemicals. We are currently working to further engineer a glycerol fermentation pathway while decreasing dependence on oxygen as well as increasing the efficiencies of fermentation and utilising the high reduction merit of glycerol for improving the fermentation efficiencies of other carbons, while looking forward use crude glycerol from biodiesel industries in the next studies.

In summary, we show here the efficient modelling of glycerol traffic to ethanol production in *S. cerevisiae*. This systematic metabolic engineering includes integration of the following: (i) imposing vigorous expression of all genes in the glycerol oxidation pathway DHA including *TPI1*; (ii) prevalence of glycerol oxidation by an oxygen-dependent dynamic by the water-forming NADH oxidase *LlNoxE*, which controls the reaction stoichiometries with regeneration of the cofactor NAD^+^; (iii) revoking the first step of both glycerol biosynthesis and glycerol catabolism through G3P (Fig. 1). Our study provides an advanced example of metabolic engineering for re-routing glycerol traffic in *S. cerevisiae* while tracking ethanol production to levels that have not yet been attained within any other safe model organisms, either native or genetically engineered (Yu *et al*, 2010; Yazdani & Gonzalez, 2008; Trinh & Srienc, 2009; Loaces *et al*, 2016; Aßkamp *et al*, 2019). Enormous global demands for bioethanol have been reported despite limited resources. Thus, this restricts global annual bioethanol production to approximately 29.1 million gallons, which represents <2.7% of transportation fuels (World bioenergy association, 2018; Renewable Fuels Association, 2018; Garside, 2020). Therefore, the present metabolic strategy represents a pivotal breakthrough in the field of utilising the glycerol surplus and provides promising scenarios for biorefinery using the glycell process. The outcome of this study promotes the association between bioethanol or bio-based chemicals and biodiesel industries, which may develop expansions without overburdening sustainability, and may also prevent a decrease in present glycerol prices as well as broaden the horizons of glycerol-producing chemicals and biofuels.

## Material and methods

### Strains, primers, cassettes, and plasmids constructions

Ancestor and all recombinant strains used in this study are listed in Table 1 and Fig. S3. The plasmids used in this study are listed in Table 6. All primers used here are listed in Table S1. Details of DNA fragments, cassettes and plasmids construction are explained as follows:

### Construction of pPGK-*ScTPI1*, *ScDAK2, ScDAK1 ScGCY1, ScSTL1* and pPGK-*ScTPI1, ScGUT2, ScGUT1, ScSTL1* plasmids

We obtained the genes from genomic DNA of the ancestor strain D452-2 to clone the plasmids in this section. Initially, cell walls were disrupted by resuspension with picked cells in 20 μl of 30 mM NaOH at 95°C for 10 min, and then used directly as a template for PCR; 1μl of fresh disrupted cells are suitable for a 50 μl of PCR mixture. All primers used to obtain the native genes were designed based on the sequences available on the Saccharomyces Genome Database (SGD): https://www.yeastgenome.org/. For assembling the plasmids pPGK-*ScTPI1*, pPGK-*ScDAK2*, pPGK-*ScDAK1*, pPGK-*ScGCY1* and pPGK-*ScSTL1*, the following genes from genomic DNA of the ancestral strain were obtained by PCR: *STL1*, *GCY1*, *DAK1*&*2* and *TPI1*. High fidelity polymerisation of KOD-plus neo with their corresponded primers (Table S1, section 1) was used during this amplification. The Xhol site of *DAK2* was deleted before cloning. Genes were purified from the PCR mixtures using columns and accessories obtained from Nippon Genetics Co., Ltd., and their cohesive ends were formed according to the designated primers and restriction enzymes. We first separately cloned each gene in the pPGK/URA3 plasmid (Kang *et al*, 1990) under the control of the expression system PGK promoter and its terminator (Table 6). We further replaced the URA3 gene in a pPGK-URA3 plasmid with an HIS3 (Rose *et al*, 1991) gene using a synthetically added overlap sequence from the pPGK plasmid to the HIS3 marker using PCR and primers (Table S1, section 3). A Gibson Assembly Master Mix was then used to assemble the overlapping ends of the two fragments to form the PGK-HIS3 plasmid. With construction of the pPGK-HIS3 plasmid (Table 6), we selected the HIS3 locus for homologous recombination in *S. cerevisiae* after linearising the plasmid at the BsiWI site. We obtained the plasmids and confirmed their gene sequences via sequencing using relevant primers (Table S1, section 2). Next, we cut the XhoI/SalI-*TPI1* cassette and inserted it into XhoI/SalI sites of a newly constructed pPGK-HIS3 plasmid. Then the genes and their systems were integrated together into one plasmid by connecting the *DAK2* set into the SalI site of the template plasmid initiated here by pPGK-*TPI1*. The non-reopened ligations (XhoI/SalI sites) were used repeatedly during the ligation of new cassettes to form new plasmids. *DAK1*, *GCY1* and *STL1* were repeatedly combined. Ultimately, the *pPGK-ScTPI1-ScDAK2-ScDAK1-ScGCY1-ScSTL1* plasmid was constructed (Table 6). Continuing with the same procedures, the plasmid *pPGK-ScTPI1-ScGUT2-ScGUT1 - ScSTL1* was also established.

### Construction of TDH3p-d22DIT1t, *TDH3-d22-OpGDH* and *TDH3-d22-LlNoxE* plasmids

#### Cassette 1: partial end of GPD1promoter-TDH3p-d22DIT1terminator-partial front side of GPD1 terminator and TDH3p-d22DIT1t plasmid

The mutated terminator d22DIT1t was purchased from Integrated DNA Technology (IDT; Tokyo, Japan) according to the published sequences (Ito *et al*, 2016). TDH3 promoter magnified from the genomic DNA of the ancestor strain D452-2 using PCR and the designated primers (Table S1, Section 4). All primers were purchased from FASMAC Company, Japan. Moreover, flanking sequences added upstream of the promoter and downstream of the terminator using the feature of PCR polymerisation with primers possessing a desired long tail, and a further extension to those flanking sequences with the addition of restriction sites was accomplished by PCR in the second step (Table S1, section 4). Then, cohesive ends of those coupled DNA fragments were processed by the restriction enzymes XhoI, NotI for the first fragment and NotI, SalI for the second. After purification of the fragments using agarose gels and columns (Nippon Genetics Co., Ltd.), one-step cloning was employed coupled the TDH3 promoter and mutated DITI terminator into XhoI/SalI of PGK/URA3 plasmid. Then the TDH3p-d22DIT1t-URA3 plasmid was constructed (Table 6).

#### Cassette 2: partial end of GPD1promoter-TDH3p-*OpGDH*-d22DIT1t-partial front side of GPD1terminator and TDH3-d22-*OpGDH* plasmid

The previously constructed TDH3p-d22DIT1t/URA3 plasmid was used as a template for construction of the next plasmid by further cloning *OpGDH*, deposited in GenBank under the accession number XP_018210953.1. Synthetic *OpGDH* was purchased from IDT. Primers are listed in Section 4 in Table S1, and full sequences are available in Table S2.

#### Cassette 3: partial end of GPD1promoter-TDH3p-*LlNoxE*-d22DIT1t-partial front side of GPD1 terminator and TDH3-d22-*LlNoxE* plasmid

We also purchased the water-forming NADH oxidase gene of *Lactococcus lactis* (IDT) based on sequence available on gene bank accession number AAK04489.1 and cloned it into TDH3p-d22DIT1t to assemble TDH3-d22-*LlNoxE* plasmid (Tables 6 and S2).

#### Cassette 4: partial end of GPD2promoter-TDH3p-*LlNoxE*-d22DIT1t-partial front side of GPD2 terminator and TDH3-d22-*LlNoxE* plasmid

In this step, we replaced the flanking sequences of the GPD1 promoter and terminator with GPD2 using PCR and the primers listed in Section 5 - of Table S1.

### Construction of multiplex pCAS-gRNA-CRISPR systems

The multiplex pCAS-gRNA system was a gift from Prof. Jamie Cate (Addgene plasmid # 60847; https://www.addgene.org/60847/) (Ryan *et al*, 2014). We used an online tool for the rational design of CRISPR/Cas target to allocate the highest probability of the on-target sites for the gRNA in the genomic DNA of *S. cerevisiae* (https://crispr.dbcls.jp/) (Naito *et al*, 2015). Accordingly, the sequences of the primers were designed based on the previously allocated sequence (20 bp before the PAM), with another 20 bp from sgRNA or HDV ribozyme for overlap (Table S1, sections 4.2, 5.1 and 7.1). First, PCR was used to synthesise two fragments from the template, the pCAS-gRNA plasmid. The first was amplified using the forward primer called pCas forward (For) located upstream of the gRNA scaffold at the SmaI site of pCas, plus the antisense primer with a reverse sequence of the target gRNA. The second fragment was amplified by forward primer with a sense sequence of gRNA and the reverse primer called pCas reverse (Rev) located downstream of the gRNA scaffold (Table S1, section 4.2). After purifying each DNA segment, overlapping and integration was carried out by PCR using the pCas For and Rev primers. The produced fragment was then restricted to the SmaI-PstI sites for cloning into a truncated pCAS-gRNA plasmid with SmaI-PstI. As a result, a new multiplex pCAS-gRNA plasmid was formed. Steps have been repeatedly performed with constructing all multiplex pCAS-gRNA plasmids targeting *ScGPD1, ScGPD2* and *GUT1* (Table 6). We confirmed the newly constructed systems by sequencing their entire scaffolds.

### Construction of pAUR101-*CuFPS1* and pAUR101-*CuFPS1*, *ScTPI1, ScDAK2, ScDAK1* plasmids

*Candida utilis* (NBRC 0988) was obtained from the National Biological Resource Center (NBRC) of National Institute of Technology and Evaluation (Japan) and was used as a template for obtaining the gene glycerol facilitator *FPS1* (*CuFPS1*). The sequence of *CuFPS1* was included in the deposited gene bank accession number BAEL01000108.1. The original pAUR101 plasmid was purchased from Takara Bio, Inc., Japan, and the primers were used to establish this plasmid listed (Table S1, section 6). A full sequence for the cassette PGK-*CuFPS1*-RPL41Bt was transferred (Table S2). First, we constructed a pAUR101-PGKp-RPL41Bt vector by one-step cloning of the SmaI-Not1 PGK promoter (fragment 1) and NotI-SalI-RPL41B terminator (fragment 2) into the SmaI-SalI pAUR101 vector and then cloning a cohesive ended *NotI-CuFPS* gene into the dephosphorylated NotI site of pAUR101-PGK-RPL41B vector to assemble pAUR101-PGKp-*CuFPS1*-RPL41Bt vector. To constitute the pAUR101-*CuFPS1, ScTPI1, ScDAK2, ScDAK1* plasmid, we detached the set of cassettes—*ScTPI1, ScDAK2* and *ScDAK1—* from previously constructed plasmids, pPGK-*ScTPI1*, *ScDAK2* and *ScDAK1* (Table S1), using restriction enzymes Xhol-SalI and reinserted that set of cassettes (*ScTPI1*, *ScDAK2* and *ScDAK1*) into the SalI site of the pAUR101-PGK-*CuFPS1*-RPL41B plasmid (Table 6).

### Construct Module M1; *CuFPS1, OgGDH, ScDAK1, ScTPI1* cassettes with flanking sequences of GUT1 promoter and terminator in plasmid pAUR101

We first obtained all fragments which could form the module M1 separately by PCR (Fig. S2); the *CuFPS1, OpGDH* genes, and mutated d22DIT terminator amplified from their synthetic DNA stocks, whereas other fragments were magnified from the genomic DNA of the D452-2 strain (Fig. S2). The full sequence of the module M1 is also accessible (Table S2), and the primer details are listed in Table S1, Section 7. Purification of the 12 amplified DNA fragments was carried out on 1%-2% agarose gel and then recovered by the FastGene Gel/PCR Extraction Kit (Nippon Genetics Co. Ltd) according to the manufacturer’s protocol. We accordingly obtained highly purified fragments before the onset of assembly using the Gibson Assembly Master Mix. We effectively joined the first three segments seamlessly, as well as for each of the next three fragments according to the manufacturer’s protocol (Gibson). We also directly amplified each set by PCR and then purified these again on an agarose gel. We repeatedly gathered the first six segments, as well as the other six fragments, and then assembled the whole module M1. We further added the SacI site upstream of the module M1 and the SmaI site downstream. These restriction sites were provided for cloning the module M1 into SacI-SmaI sites of pAUR101 vector to form pAUR 101-M1 (Table 6). Finally, we transferred the vector pAUR 101-M1 into *E. coli* as described previously and confirmed the accurate structure of M1 by sequencing the whole module M1 from pAUR-M1.

### Transformation and recombination of strains in this study

All the previous plasmids stored in *E. coli*NEB 10-beta for further uses of production of the required plasmids or cassettes, using the heat shock method according to the procedures provided with the competent cells. All plasmid extractions were performed using the QIAprep Spin Miniprep Kit following the manufacturer’s protocol. All measurements of DNA were estimated using BioSpec-nano (Shimadzu, Japan), and DNA was stored at −20°C for future use. Yeast transformation by Fast Yeast Transformation™ kit (Takara Bio) was used for integrated linear pAUR101 vector and its associated genes in the AUR1-C locus, as well as linear pPGK plasmid with its cloned genes in either HIS3 or URA3 loci (Hosaka *et al*, 1992). For achieving genome editing and the replacement of *ScGPD1, ScGPD2* and *GUT1* genes with its designated DNA repairing cassette or module, we used the protocol of CRISPR-Cas9 genome engineering in *S. cerevisiae* cells (Ryan *et al*, 2016). We confirmed target replacements using PCR for the inserted repairing cassettes with primers from upstream and downstream of the flanking recombined loci. The primers used are listed in (Table S1). Furthermore, we cultivated up to 10 generations of the selected evolved strains to confirm the loss of pCAS plasmid and re-confirm the recombination. All recombination strains and their genotypes are listed in Table 1.

### Preparation of cell-free extract

Cell proteins were extracted as previously described (Nguyen & Nevoigt, 2009; Khattab *et al*, 2013) with some modifications. The recombinant strains listed in Table 2 were cultivated in closed 50-ml Falcon tubes with 10 ml of YP medium supplemented with (w/v) 1.5% glucose and 7% glycerol (YPD_15_G_70_) for 15h with 200 rpm of shaking. Cell pellets were harvested by centrifugation at 4000 rpm for 2 min at 4 °C, then washed with 20 ml of 100 mM of HEPES buffer (pH 7.4) and centrifuged again. Then, the cell pellets were lysed in 1 ml of HEPES buffer supplemented with 1 mM MgCl2 and 10 mM 2-mercaptoethanol with approximately 400 mg of glass beads in a 2-ml Eppendorf tube. The lysis was accomplished by vigorous shaking by bench vortex with six-time intervals on ice for 30 seconds. The crude proteins were separated from the glass beads and cell debris by two rounds of centrifugation at 15000 rpm at 4°C for 5 min. Total protein concentration was estimated using Bio-RAD Quick Start Bradford 1 x dye reagent by measuring the absorbance at 595nm using BSA as a standard.

### Enzyme assays

The specific activity of glycerol dehydrogenase (*GDH*) was assayed by monitoring the increase of NADH absorbance at 340 nm within 1 min (UV-2700; UV-VIS spectrophotometer, Shimadzu, Japan) in a 1-ml mixture of 80 mM HEPES buffer (pH 7.4), 5 mM NAD^+^, 100 mM glycerol and 10 μl of crude extract. The activity of *TPI1* was assayed as described previously (Plaut & Knowles, 1972) with some modifications. The reaction mixture was 1 ml in volume and was composed of 100 mM triethanolamine hydrochloride (pH 7.53), 2.5 mM NAD^+^, 10 mM DHAP and 10 μl of crude extract. Dihydroxyacetone kinase was assayed using a universal kinase activity kit according to the manufacturer’s instructions (Catalog Number EA004).

### Intracellular concentration of NAD^+^/NADH

The cellular contents of NAD^+^/NADH were colorimetric quantitatively determined at 565 nm using a BioAssay Systems (E2ND-100) EnzyChrom™ NAD^+^/NADH Assay Kit.

### Fermentation procedures and analysis

The initial fermentation experiments were performed in 100-ml Erlenmeyer flasks with 20ml of culture medium for the estimation of micro-aerobic conditions and 20:200 liquid: flask volume for the semi-aerobic case. For the aerobic conditions, 20ml of culture medium injected into 300 or 500-ml flasks respectively, with YNB and YP medium. Three different agitation speeds were tested through this study (150, 180 and 200 rpm) at 30°C, which stated in the legends of each table or figure. For strict anaerobic condition, 20 ml of YNB medium was transferred into sterilised 50-ml vial and then caped with precision seal septa cap. Afterward, the nitrogen gas was purged into the culture medium through one use of Terumo needles to replace the dissolved oxygen and the air. Sampling was also drawn through new Terumo needles. The fermenter cells were harvested from the same volume of the pre-culture YPD medium for approximately 15h. For SK-FGG, the cultivation culture medium was YPD20G70. Cells were harvested by centrifugation at 6000 × g for 5 min at 4°C and washed with sterile water. Then, collected cells were resupplemented with the YP medium with glucose, glycerol, or both, as shown in Figures 2-3 and tables 2-5. Different initial concentrations were used to determine the fermentation abilities at those concentrations as well as with the fed-batch to estimate the maximum product under these unprecedented fermentation conditions. The cell density was monitored using spectrophotometry at 600 nm (AS ONE, Japan).

All analyses were performed using auto-sampling a 10 microliter to injected in an Aminex HPX-87H column (Bio-Rad Laboratories, Hercules, CA, USA), analysed in a refractive index detector (RID-10A; Shimadzu), and a prominence diode array detector (SPD-M20A; Shimadzu) equipped with an auto-sampling ultra-fast liquid chromatography (Shimadzu, Japan). Fractionation was accomplished at a flow rate of 0.6 ml/min with 5 mM H_2_SO_4_ as the mobile phase at 50°C. Reactant concentrations were estimated by monitoring the peak areas compared with the standards of the authenticating reactant’s glucose, glycerol, ethanol, acetic acid, pyruvate, succinic acid and acetaldehyde, which detected by RID. We used the SPD to estimate DHAP and DHA. DHA was monitored by the RID where the glycerol peaks overlapped. The measurements of DHAP were more precise when using the SPD compared when using the RID, and there was partially overlapping of the frequently negative peak of the difference of samples with the mobile phase. The detectable quantities were reported in the tables or figures and the quantities under the undetectable levels were omitted.

## Author contributions

S.M.R.K. was responsible for the research idea, conception, planning and organisation of the experiments. S.M.R.K. also provided the information for purchasing the strains, chemicals, and toolboxes for the genetic engineering, performed the experiments and analysed and discussed the results. S.M.R.K. wrote, revised, and submitted the manuscript. T.W. was responsible for all financial support and provided all chemicals and equipment. T.W. was involved in the research idea, the conception, the planning and organisation, the discussion of the results and the manuscript revision and submission.

## Data and materials availability

All necessary data required to assess our findings are available in this manuscript or its supplementary data. Further details related to this article may be requested from the authors.

## References

Ansell R, Granath K, Hohmann S, Thevelein JM, Adler L (1997) The two isoenzymes for yeast NAD^+^-dependent glycerol 3-phosphate dehydrogenase encoded by GPD1 and GPD2 have distinct roles in osmoadaptation and redox regulation. EMBO J 16(9): 2179–218

Aßkamp MR, Klein M, Nevoigt E (2019) Involvement of the external mitochondrial NADH dehydrogenase Nde1 in glycerol metabolism by wild-Type and engineered Saccharomyces cerevisiae strains. FEMS Yeast Res 1: 19(3)

Aßkamp MR, Klein M, Nevoigt E (2019) Saccharomyces cerevisiae exhibiting a modified route for uptake and catabolism of glycerol forms significant amounts of ethanol from this carbon source considered as ‘non fermentable’. Biotechnol Biofuels 12: 257

Babazadeh R, Lahtvee PJ, Adiels CB, Goksör M, Nielsen JB, Hohmann S (2017) The yeast osmostress response is carbon source dependent. Sci Rep 7: 990

Bae S-J, Kim S, Hahn J-S (2016) Efficient production of acetoin in Saccharomyces cerevisiae by disruption of 2,3-butanediol dehydrogenase and expression of NADH oxidase. Sci Rep 6: 27667

Deepak D, Gregory S (2011) Relative potential of biosynthetic pathways for biofuels and bio-based products. Nat Biotechnol 29: 1074–1078

Gancedo C, Gancedo JM, Sols A (1968) Glycerol metabolism in yeasts pathways of utilization and production. Eur J Biochem 5: 165–172

Garside M (2020) STATISTA. Fuel ethanol production worldwide in 2019, by country. https://www.statista.com/statistics/281606/ethanol-production-in-selected-countries/

Grauslund M, Lopes JM, Ronnow B (1999) Expression of GUT1, which encodes glycerol kinase in Saccharomyces cerevisiae, is controlled by the positive regulators Adr1p, Ino2p and Ino4p and the negative regulator Opi1p in a carbon source-dependent fashion. Nucleic Acids Res 27: 4391–4398

Grauslund M, Ronnow B (2000) Carbon source-dependent transcriptional regulation of the mitochondrial glycerol-3-phosphate dehydrogenase gene, GUT2, from Saccharomyces cerevisiae. Can J Microbiol 46: 1096–1100

Grüning NM, Du D, Keller MA, Luisi BF, Ralser M (2014) Inhibition of triosephosphate isomerase by phosphoenolpyruvate in the feedback-regulation of glycolysis. Open Biol 4: 130232

Ho P-W, Swinnen S, Duitama J, Nevoigt E (2017) The sole introduction of two single point mutations establishes glycerol utilization in Saccharomyces cerevisiae CEN.PK derivatives. Biotechnol Biofuels 10: 10

Hosaka K, Murakami T, Kodaki T, Nikawa J, Yamashita S (1990) Repression of choline kinase by inositol and choline in Saccharomyces cerevisiae. J Bacteriol 172(4): 2005–12

Hosaka K, Nikawa J, Kodaki T, Yamashita S (1992) A dominant mutation that alters the regulation of INO1 expression in Saccharomyces cerevisiae. J Biochem 111: 352–358

Islam Z-U, Klein M, Ødum ASR, Nevoigt E (2017) A modular metabolic engineering approach for the production of 1,2-propanediol from glycerol by Saccharomyces cerevisiae. Metab Eng 44: 223–35

Ito Y, Kitagawa T, Yamanishi M, Katahira S, Izawa S, Irie K, Furutani-Seiki M, Matsuyama T (2016) Enhancement of protein production via the strong DIT1 terminator and two RNA-binding proteins in Saccharomyces cerevisiae, Sci. Rep 6: 36997

Ito Y, Yamanishi M, Ikeuchi A, Imamura C, Tokuhiro K, Kitagawa T, Matsuyama T (2013) Characterization of five terminator regions that increase the protein yield of a transgene in Saccharomyces cerevisiae. J Biotechnol 168: 486–492

Kampf G, Todt D, Pfaender S, Steinmann E (2020) Persistence of coronaviruses on inanimate surfaces and their inactivation with biocidal agents. J Hosp Infect 104: 246–251

Kang YS, Kane J, Kurjan J, Stadel JM, Tipper DJ (1990) Effects of expression of mammalian G alpha and hybrid mammalian-yeast G alpha proteins on the yeast pheromone response signal transduction pathway. Mol Cell Biol 10: 2582–2590

Kato M, Lin SJ (2014) Regulation of NAD^+^ metabolism, signaling and compartmentalization in the yeast Saccharomyces cerevisiae. DNA Repair 23: 49–58

Khattab SMR, Kodaki T (2014) Efficient bioethanol production by overexpression of endogenous Saccharomyces cerevisiae xylulokinase and NADPH-dependent aldose reductase with mutated strictly NADP^+^-dependent Pichia stipitis xylitol dehydrogenase. Process Biochem 49: 1838–1842

Khattab SMR, Saimura M, Kodaki T (2013) Boost in bioethanol production using recombinant Saccharomyces cerevisiae with mutated strictly NADPH-dependent xylose reductase and NADP+-dependent xylitol dehydrogenase. J Biotechnol 165: 153–156

Khattab SMR, Watanabe T (2019) Bioethanol from sugarcane bagasse: Status and perspectives. In Bioethanol Production from Food Crops: Sustainable Sources, Interventions, and Challenges, Ramesh CR, Ramachandran S (eds) pp 187–212. Elsevier

Kim J-W, Lee Y-G, Kim S-J, Jin, Y-S, Seo, J-H. (2019) Deletion of glycerol-3-phosphate dehydrogenase genes improved 2,3-butanediol production by reducing glycerol production in pyruvate decarboxylase-deficient Saccharomyces cerevisiae. J Biotechnol 304: 31–37

Kim J-W, Seo S-O, Zhang G-C, Jin Y-S, Seo J-H (2015) Expression of Lactococcus lactis NADH oxidase increases 2,3-butanediol production in Pdc-deficient Saccharomyces cerevisiae. Bioresour Technol 191: 512–519

Kim S, Hahn J-S. (2015) Efficient production of 2,3-butanediol in Saccharomyces cerevisiae by eliminating ethanol and glycerol production and redox rebalancing. Metab Eng 31: 94–101

Klein M, Carrillo M, Xiberras J, Islam Z-U, Swinnen S, Nevoigt E (2016) Towards the exploitation of glycerol’s high reducing power in Saccharomyces cerevisiae-based bioprocesses. Metab Eng 38: 464–472

Klein M, Islam Z-U, Knudsen PB, Carrillo M, Swinnen S, Workman M, Nevoigt E (2016) The expression of glycerol facilitators from various yeast species improves growth on glycerol of Saccharomyces cerevisiae. Metab Eng Commun 3: 252–257

Kodaki T, Yamashita S (1989) Characterization of the methyltransferases in the yeast phosphatidylethanolamine methylation pathway by selective gene disruption. Eur J Biochcm 185: 243–251

Larsson C, Påhlman IL, Ansell R, Rigoulet M, Adler L, Gustafsson L (1998) The importance of the glycerol 3-phosphate shuttle during aerobic growth of Saccharomyces cerevisiae. Yeast 15;14(4): 347–357

Loaces I, Rodríguez C, Amarelle V, Fabiano E, Noya F (2016) Improved glycerol to ethanol conversion by E. coli using a metagenomic fragment isolated from an anaerobic reactor. J Ind Microbiol Biotechnol 43: 1405–1416

Lovins, AB, Datta EK, Bustnes O-E, Koomey JG & Glasgow NJ (2004) Winning the Oil Endgame. Innovation for Profits, Jobs, and Security. Snowmass, CO: Rocky Mountain Institute, United States

Luque R, Herrero-Davila L, Campelo JM, Clark JH, Hidalgo JM, Luna D, Marinas JM, Romero AA (2008) Biofuels: a technological perspective. Energy Environ Sci 1: 542–564

Lutstorf U, Megnet R (1968) Multiple forms of alcohol dehydrogenase in Saccharomyces cerevisiae. physiological control of ADH-2 and properties of ADH-2 and ADH-4. Arch Biochem Biophys 126: 933–944

Meadows AL, Hawkins KM, Tsegaye Y, Antipov E, Kim Y, Raetz L, Dahl RH, Tai A, Mahatdejkul-Meadows T, Xu L et al (2016) Rewriting yeast central carbon metabolism for industrial isoprenoid production. Nature 537: 694–697

Molin M, Norbeck J, Blomberg A (2003) Dihydroxyacetone kinases in Saccharomyces cerevisiae are involved in detoxification of dihydroxyacetone. J Biol Chem 278(3): 1415–1423

Murata K, Fukuda Y, Watanabe K, Saikusa T, Shimosaka M, Kimura A (1985) Characterization of methylglyoxal synthase in Saccharomyces cerevisiae. Biochem Biophys Res Commun 131: 190–198

Naito Y, Hino K, Bono H, Ui-Tei K (2015) CRISPRdirect: software for designing CRISPR/Cas guide RNA with reduced off-target sites. J Bioinform 31: 1120–1123

Nambu-Nishida Y, Sakihama Y, Ishii J, Hasunuma T, Kondo A (2018) Selection of yeast Saccharomyces cerevisiae promoters available for xylose cultivation and fermentation. J Biosci Bioeng 125(1): 76–86

Nguyen HT, Nevoigt E (2009) Engineering of Saccharomyces cerevisiae for the production of dihydroxyacetone (DHA) from sugars: A proof of concept. Metab Eng 11: 335–346

Nomanbhay S, Hussein R, Ong MY (2018) Sustainability of biodiesel production in Malaysia by production of bio-oil from crude glycerol using microwave pyrolysis: a review. Green Chem Lett Rev 11: 135–157

Ochoa-Estopier A, Lesage J, Gorret N, Guillouet SE (2011) Kinetic analysis of a Saccharomyces cerevisiae strain adapted for improved growth on glycerol: implications for the development of yeast bioprocesses on glycerol. Bioresour Technol 102(2): 1521–7

Ohashi Y, Watanabe T (2018) Catalytic performance of food Additives Alum, flocculating agent, Al(SO4)3, AlCl3 and other Lewis acids in microwave solvolysis of hardwoods and recalcitrant softwood for biorefinery. ACS Omega 3:16271–16280

Overkamp KM, Bakker BM, Kötter P, Luttik MAH, van Dijken JP, Pronk, JT (2002) Metabolic Engineering of glycerol production in Saccharomyces cerevisiae. Appl Environ Microbiol 68(6): 2814–2821

Panozzo C, Nawara M, Suski C, Kucharczyka R, Skoneczny M, Bécam A M, Rytka J, Herbert, CJ (2002) Aerobic and anaerobic NAD^+^ metabolism in Saccharomyces cerevisiae. FEBSLett 517(1-3): 97–102

Peris D, Moriarty RV, Alexander WG, Baker EC, Sylvester K, Sardi M, Langdon QK, Libkind D, Wang Q-M, Bai F-Y et al. (2017) Hybridization and adaptive evolution of diverse Saccharomyces species for cellulosic biofuel production. Biotechnol Biofuels 10: 78

Plaut B, Knowles J R (1972) pH-dependence of the triose phosphate isomerase reaction. Biochem J 129(2): 311–320

Ragauskas AJ, Williams CK, Davison BH, Britovsek G, Cairney J, Eckert CA, Frederick-Jr WJ, Hallett JP, Leak DJ, et al (2006) The path forward for biofuels and biomaterials. Science 311: 484–489

Renewable Fuels Association. World Fuel Ethanol Production 2018-. https://ethanolrfa.org/statistics/annual-ethanol-production/

Roberts GG, Hudson AP (2009) Rsf1p is required for an efficient metabolic shift from fermentative to glycerol-based respiratory growth in S. cerevisiae. Yeast 26: 95–110

Roberts TM, Kaltenbach H-M, Rudolf F (2020) Development and optimisation of a defined high cell density yeast medium. Yeast 37(5-6): 336–347

Rose MD, Broach JR (1991) Cloning genes by complementation in yeast. Methods Enzymol 194: 195–230

Ryan OW, Poddar S, Cate JHD. (2016) CRISPR-Cas9 Genome engineering in Saccharomyces cerevisiae cells. Cold Spring Harbor Protocol

Ryan OW, Skerker JM, Maurer MJ, Li X, Tsai CJ, Poddar S, Lee ME, DeLoache W, Dueber JE, Arkin AP, Cate JHD (2014) Selection of chromosomal DNA libraries using a multiplex CRISPR system. eLife 3: e03703

Saito H, Posas F (2012) Response to Hyperosmotic Stress. Genetics 192: 289–318

Semkiv M, Kata I, Ternavska O, Sibirny W, Dmytruk K, Sibirny A (2019) Overexpression of the genes of glycerol catabolism and glycerol facilitator improves glycerol conversion to ethanol in the methylotrophic thermotolerant yeast Ogataea polymorpha. Yeast 36: 329–339

Shi Y, Vaden DL, Ju S, Ding D, Geiger JH, Greenberg ML (2005) Genetic perturbation of glycolysis results in inhibition of de novo inositol biosynthesis. J Biol Chem280(51): 41805–41810

Sprague GF, Cronan JE (1977) Isolation and characterization of Saccharomyces cerevisiae mutants defective in glycerol catabolism. J Bacteriol 129(3):1335–42

Swinnen S, Ho P-W, Klein M, Nevoigt E (2016) Genetic determinants for enhanced glycerol growth of Saccharomyces cerevisiae. Metab Eng 36: 68–79

Swinnen S, Klein M, Carrillo M, McInnes J, Nguyen HT, Nevoigt E (2013) Reevaluation of glycerol utilization in Saccharomyces cerevisiae: characterization of an isolate that grows on glycerol without supporting supplements. Biotechnol Biofuels 6: 157

Trinh CT, Srienc F (2009) Metabolic engineering of Escherichia coli for efficient conversion of glycerol to ethanol. Appl Environ Microbiol 75: 6696–6705

Turcotte B, Liang XB, Robert F, Soontorngun N (2010) Transcriptional regulation of nonfermentable carbon utilization in budding yeast. FEMS Yeast Res 10: 2–13

Vemuri GN, Eiteman MA, McEwen JE, Olsson L, Nielsen J (2007) Increasing NADH oxidation reduces overflow metabolism in Saccharomyces cerevisiae. Proc Natl Acad Sci 104: 2402–2407

Wei L, Wang Z, Zhang G, Ye B (2017) Characterization of terminators in saccharomyces cerevisiae and an exploration of factors affecting their strength. Chem Bio Chem 18: 2422–2427

Wei N, Quarterman J, Kim SR, Cate JH. Jin YS (2013) Enhanced biofuel production through coupled acetic acid and xylose consumption by engineered yeast. Nat Commun 4: 2580.

World bioenergy association. WBA Global Bioenergy Statistics 2018 https://worldbioenergy.org/uploads/181203%20WBA%20GBS%202018_hq.pdf

Xiberras J, Klein M, Nevoigt E (2019) Glycerol as a substrate for Saccharomyces cerevisiae based bioprocesses - Knowledge gaps regarding the central carbon catabolism of this ‘non-fermentable’ carbon source. Biotechnol Adv 37: 107378

Yamada-Onodera K, Yamamoto H, Emoto E, Kawahara N, Tani Y (2002) Charaterisation of glycerol dehydrogenase from a methylotrophic yeast Hansenula polymorpha DL-1, and its gene cloning. Acta Biotechnol 22: 337–353

Yamanishi M, Ito Y, Kintaka R, Imamura C, Katahira S, Ikeuchi A, Moriya H, Matsuyama T (2013) A genome-wide activity assessment of terminator regions in Saccharomyces cerevisiae provides a “terminatome” toolbox. ACS Synth Biol 2: 337–347

Yazdani SS, Gonzalez R (2007) Anaerobic fermentation of glycerol: a path to economic viability for the biofuels industry. Curr Opin Biotechnol 18: 213–219

Yazdani SS, Gonzalez R (2008). Engineering Escherichia coli for the efficient conversion of glycerol to ethanol and co-products. Metab Eng 10: 340–351

Yu KO, Kim SW, Han SO (2010) Engineering of glycerol utilization pathway for ethanol production by Saccharomyces cerevisiae. Bioresour Technol 101: 4157–4161

Zheng Z, Zou J (2001) The initial step of the glycerolipid pathway: identification of glycerol 3-phosphate/dihydroxyacetone phosphate dual substrate acyltransferases in Saccharomyces cerevisiae. J Biol Chem 276: 41710–41716

